# Telomere dynamics parameters inferred from mathematical modeling of telomere length distribution

**DOI:** 10.64898/2026.10.15.759289

**Authors:** Viviana Gavilanes Guerrero, Marie Doumic, Zhou Xu

## Abstract

Telomeres, repeated sequences located at the ends of linear chromosomes, are essential for genome integrity. As these regions are subject to dynamics of shortening and lengthening by telomerase, they exhibit length heterogeneity. Telomere length is thus best represented by an entire distribution rather than by a single value. Long-read sequencing technologies now allow us to experimentally access this distribution at high resolution. Our working hypothesis posits that this distribution encodes information that relates to the biochemical mechanisms underlying telomere length dynamics. Here, to numerically infer the biochemical parameters that control telomere shortening and elongation by telomerase, we developed a mathematical model that we fit on long-read-sequencing-derived telomere length distributions of *Saccharomyces cerevisiae*. We prove that in our model, telomere length can be mathematically represented as an ergodic Markov chain and we provide an explicit expression of the steady-state distribution. We show that the biochemical parameters can be uniquely inferred from the distribution, which we verify on synthetic data with known ground truth parameters. Using experimental telomere length distributions of wild-type and mutants of telomere regulators, we find that our pipeline of parameter inference yields biologically relevant values. We then fit the model to the telomere length distribution of 66 natural yeast strains to reveal the diversity of telomere length regulation parameters. Furthermore, we demonstrate that individual chromosome extremities can sometimes be associated with distinct regulation parameters. Overall, our mathematical model and computational pipeline represent a powerful approach to estimate biochemical parameters affecting telomere length regulation.

## Introduction

Linear chromosomes end with telomeres, repetitive sequences essential for genome integrity and cell proliferation [1–2]. Telomeres are bound by specific proteins that prevent the detection of the chromosome ends as double-strand breaks, which would otherwise trigger the activation of the DNA damage checkpoint and inappropriate repair mechanisms, such as chromosome fusions. In the budding yeast *Saccharomyces cerevisiae*, these include Rap1 and Rif1/Rif2, which are bound to double-stranded DNA (dsDNA) in numbers roughly proportional to telomere length [3–4]. At each round of DNA replication, telomeres shorten because of the end-replication problem, the inability of the replication machinery to fully synthesize DNA up to the very end of the linear template molecule [5–7]. However, when telomeres become too short, the end protection function conferred by the telomeric proteins is partially comprised, resulting in DNA damage checkpoint activation and replicative senescence, a permanent cell cycle arrest preventing cell proliferation [8–10].

A minimal length must thus be maintained to allow for cell division. In most species, including budding yeast, this is accomplished by expressing the telomerase enzyme, which preferentially adds a variable number of telomeric repeats to shorter telomeres [8, 11–15]. As cells divide, individual telomeres are thus subject to the opposing mechanisms of shortening and preferential elongation of short telomeres, and their length reaches a broad steady-state distribution, rather than a single equilibrium value [16]. Therefore, the steady-state distribution results from the interaction of the molecular and biochemical parameters controlling the shortening mechanism, the telomerase recruitment probability, and the number of nucleotides added by telomerase. Despite the distribution inherently containing information regarding these parameters, whether it might be possible to infer these parameters from the distribution is unclear. Recently, Oxford Nanopore Technologies (ONT) long-read sequencing technologies enabled the measurement of telomere length distributions at unprecedented resolution [17–19], providing high-quality numerical data on which such a parametric inference approach might be attempted.

Mathematical modeling approaches have been applied to understand the link between telomerase activity and telomere length distribution or average telomere length at equilibrium, and to probe the consequences of length-dependent telomerase regulation for the shape of the distribution [16, 20–28]. However, previous approaches have not attempted to relate the steady-state distribution back to the biochemical parameters controlling length regulation.

Here, we propose a mathematical model of telomere length regulation, building on the stochastic framework introduced in [16] and related previous work on telomere dynamics [26]. We mathe-matically formulate the steady-state distribution as the limit of the time-average of the Markov chain defining telomere dynamics and compute an expression of the distribution, without relying on simulations. We then show that fitting the model to synthetic and experimental data allows the inference of the parameters controlling telomere length regulation, providing insights into the mechanisms of telomere shortening and elongation.

## Results

### Mathematical model of telomere length as a Markov chain

To model telomere length dynamics, we consider the discrete-time stochastic model introduced by [16]. The length of a single telomere at generation *n* is denoted by *L*_*n*_ ∈ ℕ_0_ and evolves according to

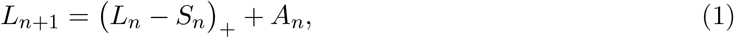

where (*x*)_+_ := max(*x*, 0) and *S*_*n*_ *>* 0 represents telomere shortening in base pairs (bp) and *A*_*n*_ ≥ 0 corresponds to telomere elongation.

The elongation term is further decomposed as *A*_*n*_ = *C*_*n*_*Y*_*n*_, where *C*_*n*_ ∈ {0, 1} is the telomerase recruitment indicator and *Y*_*n*_ ≥ 1 is the length (in bp) of elongation. Conditionally on *L*_*n*_ = *l*, these terms satisfy ℙ(*C*_*n*_ = 1 | *L*_*n*_ = *l*) = *f* (*l*), ℙ(*Y*_*n*_ = *k* | *L*_*n*_ = *l*) = *q*_*l*_(*k*), *k* ∈ ℕ*, where *f* :ℕ_0_ → [0, 1] is the recruitment function and *q*_*l*_ is the conditional distribution of the elongation length for a telomere of length *l*. The index *n* on *S*_*n*_, *C*_*n*_, and *Y*_*n*_ denotes the random quantities used during the transition from generation *n* to generation *n* + 1. At each generation, conditionally on the current length *L*_*n*_, these quantities are generated according to the same mechanism and are independent of the earlier history. In particular, the distribution of the recruitment variable *C*_*n*_ depends on the current telomere length *L*_*n*_, since *C*_*n*_ | *L*_*n*_ ~ Bernoulli(*f* (*L*_*n*_)). Under these conditions, the process (*L*_*n*_)_n*≥*0_ is a time-homogeneous Markov chain. The Markov property and the resulting transition probabilities are derived in Supp. Text, Section S1.

This general framework encodes three separate mechanistic steps affecting telomere length: *S*_*n*_, telomere shortening at each round of replication; *C*_*n*_, telomere length-dependent telomerase recruitment; and *Y*_*n*_, the number of nucleotides added by telomerase.

In this work, for the purpose of parameter inference, we specify deterministic shortening with *S*_*n*_ = *a*, for a constant integer *a* ∈ ℕ* (Fig. 1A). As in Xu et al. [16], we assume that *Y*_*n*_ follows a geometric distribution independent of *L*_*n*_: ℙ(*Y*_*n*_ = *k*) = *p*(1 − *p*)^k*−*1^, with *k* ∈ ℕ*, where *p* represents the probability that telomerase stops elongation after adding one unit of length Consequently, 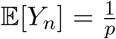 and 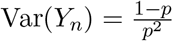. The recruitment function is written as

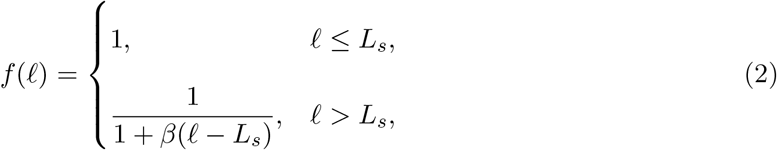

with parameters *L*_*s*_ ≥ 0, corresponding to the threshold length below which telomerase is recruited with probability 1, and *β >* 0, representing the rate at which the recruitment probability decreases above *L*_*s*_ (Fig. 1B). The formulation of *f*, *L*_*s*_, and *β* is consistent with telomerase being preferentially targeted to shorter telomeres [14, 16].

**Figure 1.**
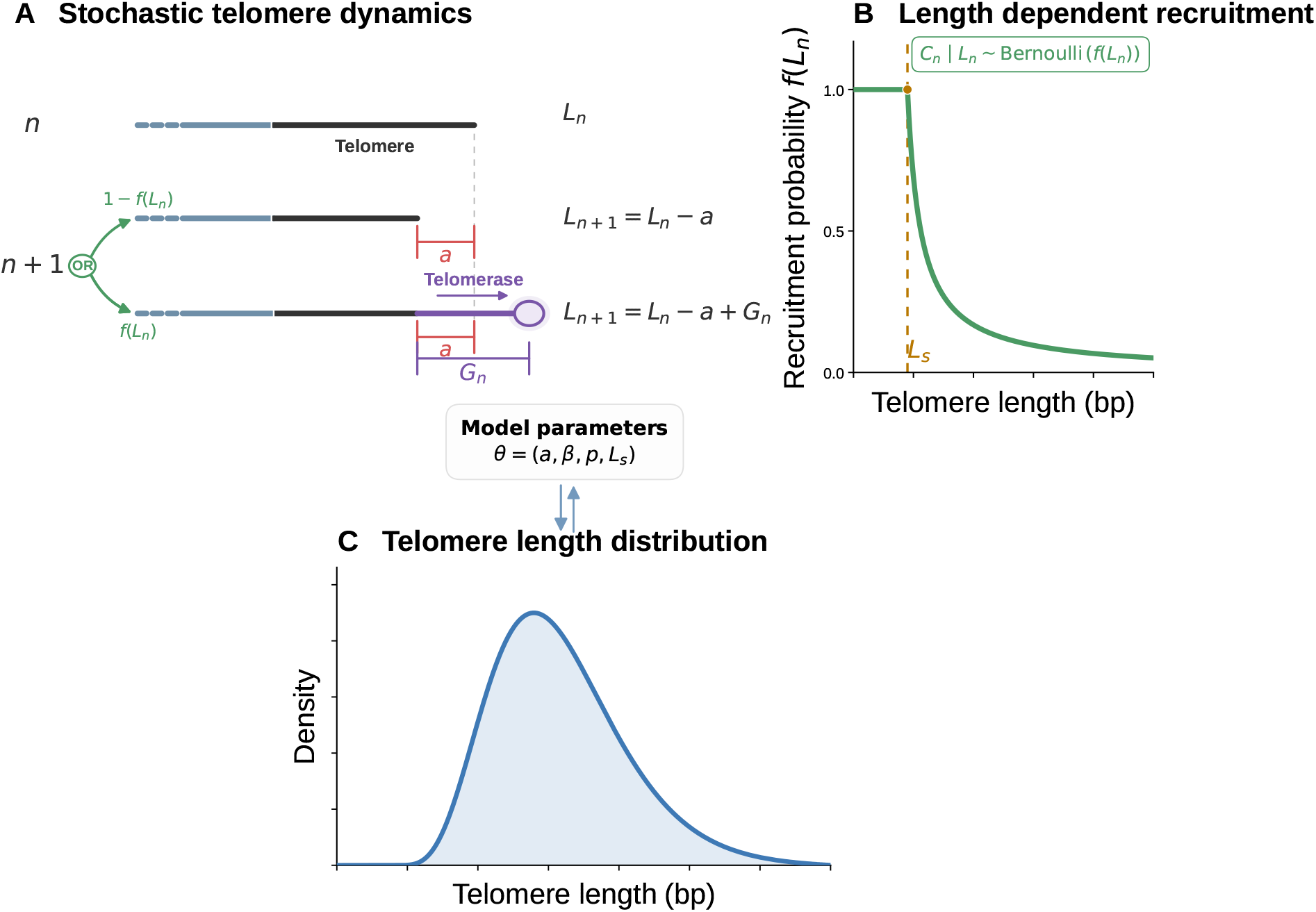
Mathematical model of telomere length dynamics. **(A)** Schematic representation of a telomere (black line) and the adjacent subtelomere (blue). Starting from a length *L*_*n*_ at generation *n*, at the next generation *n* + 1, *L*_n+1_ shortens by a length *a*. It is then either not elongated, with probability 1−*f* (*L*_*n*_), or elongated by telomerase by a random length *G*_*n*_, with probability *f* (*L*_*n*_). **(B)** The probability of telomerase recruitment *f* (*L*_*n*_) depends on telomere length: it is maximal, equal to 1, below the threshold *L*_*s*_ and decreases for longer telomeres in a manner dependent on *β*. **(C)** These mechanisms determine the steady-state telomere length distribution generated by the model, which depends on the parameters ***θ*** = (*a, β, p, L*_*s*_).

Under these assumptions, Eq. (1) becomes

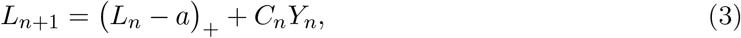

where *C*_*n*_ | *L*_*n*_ ~ Bernoulli *f* (*L*_*n*_) and *Y*_*n*_ ~ Geom(*p*). Here *C*_*n*_ and *Y*_*n*_ are the telomerase-recruitment and elongation variables associated with the update *L*_*n*_ → *L*_*n*+1_.The parameter vector for this model is *θ* = (*a, L*_*s*_, *β, p*) ∈ ℕ* ×ℕ_0_ × (0, ∞) × (0, 1). Since *θ* encapsulates the main parameters controlling telomere length, the inverse problem of inferring *θ* from experimental telomere length distributions is a powerful approach to gain insights into telomere length dynamics and regulation (Fig. 1A and C).

### The time-average of telomere length converges to a unique stationary distribution

We consider that in a sufficiently large telomerase-positive cell population, the aggregated distribution of telomere lengths is stationary and approximates the time-average of the process (*L*_*n*_)_*n≥*0_, under the assumption that telomeres behave independently of each other [29].

Let *r*_*i*_ = (*i* − *a*)_+_, the transition kernel *P* between two states *i* and *j*, i.e. two telomere lengths, is

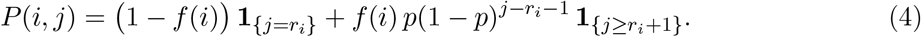

For *i* ≥ *a, r*_*i*_ = *i* − *a*; for *i < a*, telomere shortening is absorbed at 0 and therefore *r*_*i*_ = 0.

By classical results from Markov chain theory, the empirical distribution of (*L*_*n*_)_*n≥*0_ converges to a unique stationary distribution *π*_*θ*_ (Supp. Text), satisfying

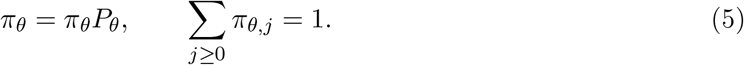

While the existence and uniqueness of the solution to this system can be established theoretically under specific conditions on *f*, *a*, and *p* [30–31], the numerical computation of *π*_*θ*_ requires a finite-dimensional approximation. We thus truncate the chain to the finite state space 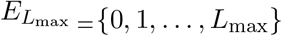, where *L*_max_ is chosen large enough so that *π*_*θ,j*_ is negligibly small for *j > L*_max_.

We prove that, for any parameter vector *θ*, the truncated transition matrix *P*_*θ*_ on 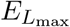 is irreducible and aperiodic (Supp. Text). As a consequence, the truncated Markov chain admits a unique stationary distribution *π*_*θ*_ satisfying Eq. (5), and the chain converges to *π*_*θ*_ starting from any initial distribution [30] (Fig. 2A and B).

**Figure 2.**
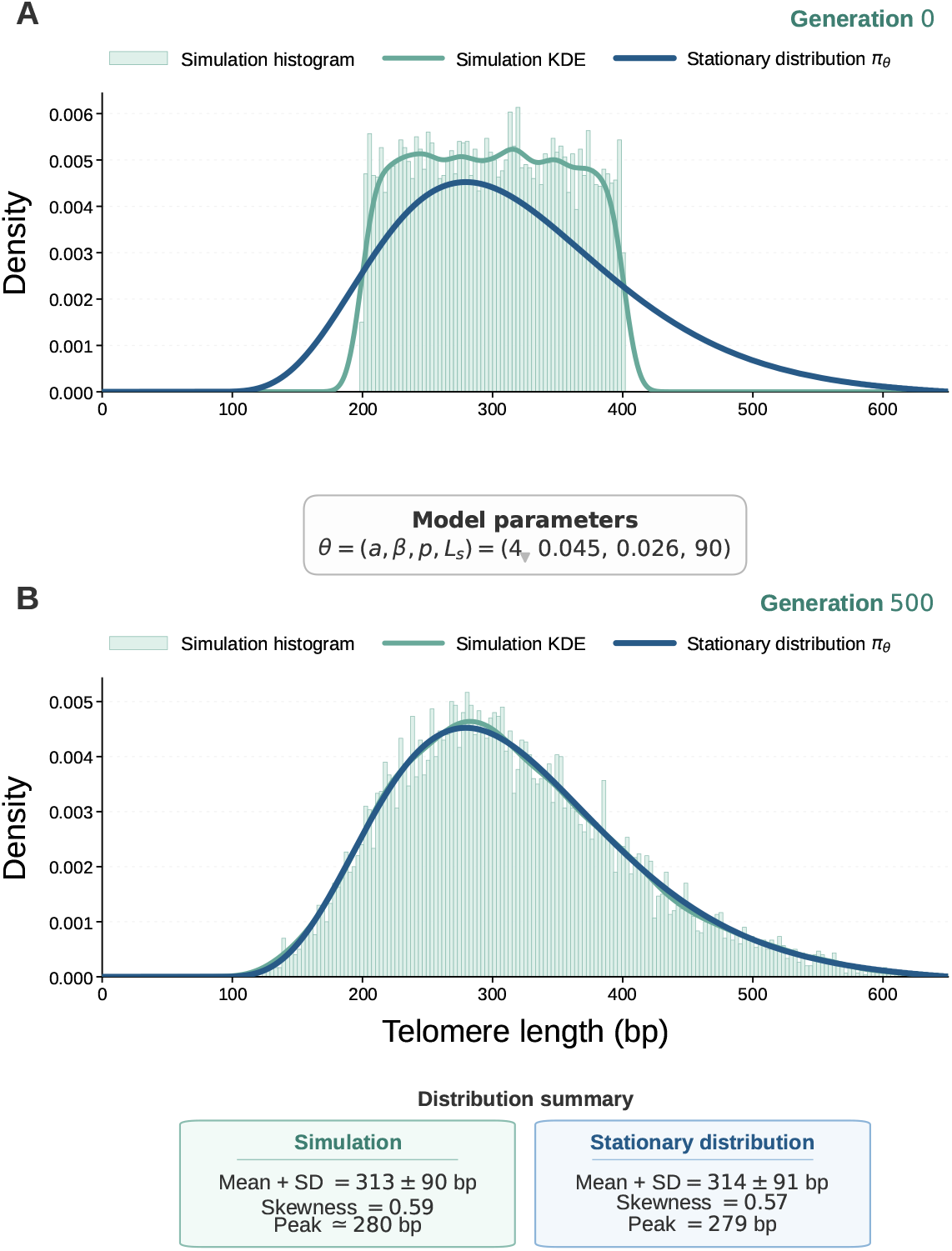
Convergence to the stationary telomere length distribution. **(A)** A set of *N* = 10,000 independent telomere lengths was initialized from a discrete uniform distribution on [200, 400] bp. The histogram and the kernel density estimation (KDE) represent the simulated lengths at generation 0, while the dark blue curve denotes the stationary distribution *π*_*θ*_ for *θ* = (*a, β, p, L*_*s*_) = (4, 0.045, 0.026, 90). **(B)** After 500 generations, the simulated distribution closely overlaps with *π*_*θ*_; their mean, standard deviation, skewness, and peak are reported below the panel.

We further show that the convergence is geometric. Specifically, if *λ*_2_ denotes the second-largest modulus among all eigenvalues of *P*_*θ*_, then for any initial distribution *v* and any *n* ≥ 0,

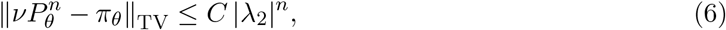

where ∥ · ∥_TV_ denotes the total variation distance and *C* is a constant that depends on *v* [30–31]. The spectral gap 1 − |*λ*_2_| *>* 0 quantifies how rapidly the chain mixes.

Thus, instead of running numerical simulations of the Markov chain, the stationary distribution will be directly computed to fit experimental telomere length distributions (Fig. 2B).

### Telomere shortening and elongation parameters can be inferred uniquely from the stationary distribution

We hypothesize that the stationary telomere length distribution contains information on telomere shortening and elongation, which could be inferred by fitting our mathematical model. This can be formulated as the following inverse problem:

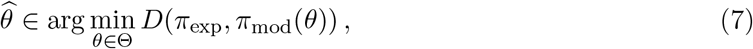

where Θ is the admissible parameter space, *π*_exp_ and *π*_mod_(*θ*) denote the empirical telomere length distribution and the stationary distribution predicted by the model with parameter *θ*, respectively, and *D* is a discrepancy measure between probability distributions. We choose the 1-Wasserstein distance (hereafter called Wasserstein distance) [32–33] because it accounts for the global redistribution of probability mass between the distributions, thus making it sensitive to differences that are spread over the support.

We first ask whether the stationary distribution uniquely determines the model parameters, that is, whether the model is structurally identifiable [34–36]. We consider this question conditional on the true value of *a* being known. We prove the following result (Supp. Text).

#### Theorem 1.

*Assume that the true shortening parameter a is known. Under the regularity conditions stated in the Supp. Text, the exact stationary distribution uniquely determines the remaining parameters* (*L*_*s*_, *β, p*).

Assuming telomere length dynamics are governed by our model, Theorem 1 therefore indicates that, when *a* is known, the stationary telomere length distribution contains sufficient information to uniquely determine the parameters controlling telomerase recruitment and elongation.

However, sampling noise introduces an irreducible variability that limits the direct application of Theorem 1 to finite experimental datasets. Even if the model were exactly correct, i.e. *π*_mod_(*θ*\*) = *π*\* for some true parameter vector *θ*\*, the empirical distribution *π*_*n*_ computed from a finite sample of size *n* would still differ from *π*\* because of sampling variability. In terms of the Wasserstein distance, this discrepancy has a natural scale of order *n*^*−*1/2^. More precisely, we obtain the following result.

#### Proposition 1

(Expected empirical Wasserstein fluctuation). *Let X*_1_, …, *X*_*n*_ *be i*.*i*.*d. with distribution π on* 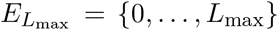, *and let* 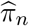 *be the corresponding empirical distribution*.

*If*

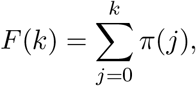

*Then*

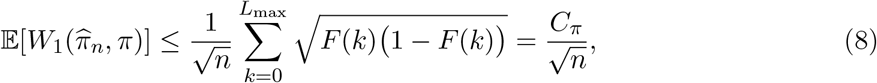

*Where*

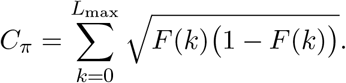

The proof is presented in the Supp. Text. Monte Carlo simulations numerically confirm the *n*^*−*1/2^ scaling (Supp. Fig. S1).

Proposition 1 provides a natural reference scale for practical identifiability. If

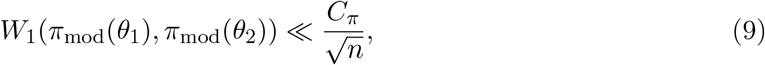

then the difference between the two predicted stationary distributions is small compared with the characteristic sampling fluctuation at sample size *n*. Although the two parameter vectors remain distinct in the structural-identifiability sense, they may therefore be *practically indistinguishable* from a finite sample [36].

We thus distinguish structural identifiability, which concerns uniqueness from the exact stationary law, from practical identifiability, which depends on the amount of experimental data available. To account for sampling variability in the empirical telomere length distributions, we associate each dataset with a bootstrap-based Wasserstein threshold *W*_thr_. Since the underlying distribution is unknown, the observed empirical distribution *π*_exp_ is used as a proxy for estimating this variability by bootstrap resampling.

To assess whether the discrepancy between the fitted stationary distribution and the experimental distribution is compatible with this sampling variability, we compare

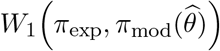

with the bootstrap-based threshold *W*_thr_ (see Section Bootstrap based Wasserstein threshold for its definition).

A fitted distance below this threshold is of the same order as variability generated by finite sampling of the observed distribution, whereas a larger distance indicates a discrepancy exceeding this estimated sampling variability.

### Benchmarking the parameter inference method on synthetic datasets

To evaluate to which extent distinct parameter vectors can be recovered from an empirical distribution, we performed simulations based on our model to generate synthetic data of sample size *N* with known ground truth parameters, and fitted these data. We chose to use the covariance matrix adaptation evolution strategy (CMA-ES) method [37] to minimize the objective function

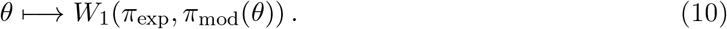

CMA-ES is a derivative-free, population-based optimizer designed for nonlinear, non-convex and ill-conditioned problems on continuous domains, and is therefore well suited for our problem (Supp. Text).

We generated synthetic data for three sets of parameters, set A, B and C, yielding realistic telomere length distributions with means at 70, 327, 609 bp respectivly, and with sample sizes *N* ranging from 100 to 10,000 for each parameter set.

To reduce the number of free parameters that need to be estimated, and because *a* is a small discrete parameter, we did not include *a* directly in the continuous CMA-ES optimization. Instead, for each candidate value *a* ∈ {2, …, 20}, we fixed *a* and performed 15 independent CMA-ES optimizations of the remaining parameters (*L*_*s*_, *β, p*). The candidate value of *a* providing the best fit, as measured by the smallest Wasserstein distance, was then selected.

In all conditions, starting from random positions in the parameter space, CMA-ES optimization moved the estimated parameters toward regions that better reproduced the synthetic data (Fig. 3A– C and Supp. Fig. S2A–F). Fig. 3A–C illustrates this behavior for set B at *N* = 3000: among the fits compatible with the bootstrap Wasserstein threshold, the final values of *L*_*s*_, *β* and *p* are considerably more concentrated than their initial values, with the true parameter values lying within or close to the region explored by the final estimates.

**Figure 3.**
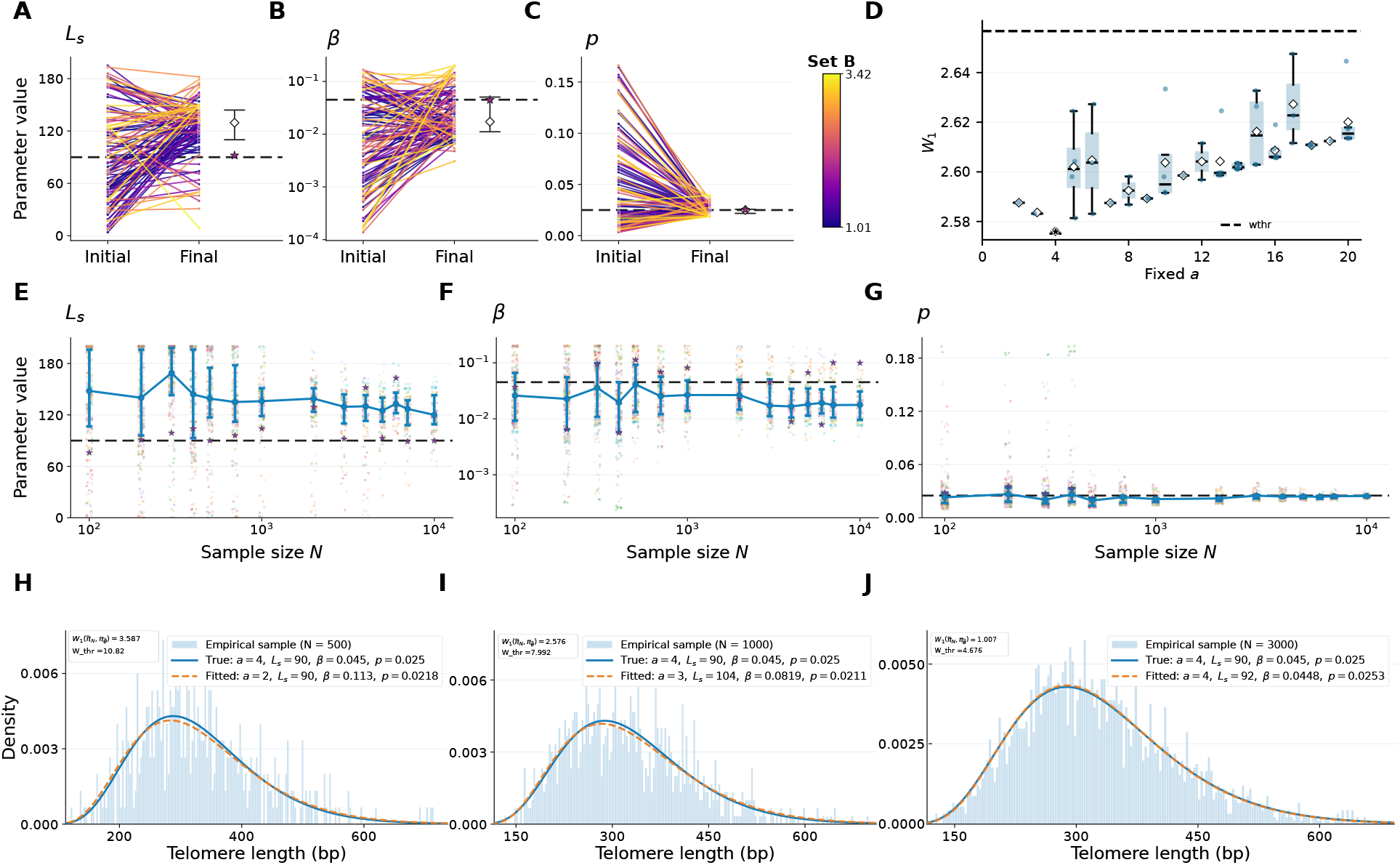
Parameter inference performance using synthetic data. Synthetic data were generated for set B with ground truth parameters (*a, L*_*s*_, *β, p*) = (4, 90, 0.045, 0.025). **(A–C)** Initial (left side) and final (right side) values of *L*_*s*_, *β*, and *p*, respectively, for fits retained at sample size *N* = 3000. Lines connect the initial and final values of individual optimization runs. Dashed lines indicate the true parameter values, while diamonds and interval bars show the median and interquartile range of the final estimates. **(D)** Wasserstein distance *W*_1_ as a function of the fixed parameter *a*. Boxplots summarize the variability of *W*_1_ across optimization runs, with diamonds indicating the median and boxes representing the interquartile range. The horizontal dashed line indicates the bootstrap-based acceptance threshold *W*_thr_ **(E–G)** Recovery of *L*_*s*_, *β*, and *p*, respectively, as a function of sample size *N*. Dashed horizontal lines indicate the ground truth values. Each colored dot represents one accepted CMA-ES optimization run. The blue dots, lines, and intervals represent the median and interquartile range across CMA-ES runs. The best fit is indicated by a star. **(H–J)** Simulated synthetic data (histogram), true telomere length distribution (blue solid line), and optimally fitted telomere length distribution (orange dashed line) for *N* = 500, *N* = 1000, and *N* = 3000, respectively. The Wasserstein distance of the optimal fit and the corresponding bootstrap threshold are reported in each panel.

We also examined the recovery of the shortening parameter *a* by fixing its value between 2 and 20 and optimizing the remaining parameters (Fig. 3D). For set B at *N* = 3000, the lowest Wasserstein distance was obtained at *a* = 4, corresponding to the ground truth value. Although the Wasserstein distance generally increased for larger values of *a*, several alternative values still produced fits below the bootstrap acceptance threshold. Thus, the optimization correctly favored the true shortening parameter while allowing satisfactory fits for other values.

The quality of parameter recovery nevertheless depended on sample size (Fig. 3E–G and Supp. Fig. S2G–L). At small sample sizes, several parameter combinations could generate stationary distributions compatible with the same empirical sample, leading to substantial variability in the recovered parameters. As *N* increased, the best fitting estimates became progressively closer to the ground truth values and their variability generally decreased. This improvement was particularly clear for the elongation parameter *p*, whereas *L*_*s*_ and *β* retained greater variability, reflecting the different practical sensitivity of the stationary distribution to the model parameters.

The fitted distributions provide a direct illustration of this finite-sample effect (Fig. 3G–I). For set B, the true parameters are (*a, L*_*s*_, *β, p*) = (4, 90, 0.045, 0.025). At *N* = 500, the optimal fitted distribution already reproduces the overall shape of the empirical distribution and its Wasserstein distance is below the bootstrap threshold, although the selected value of *a* and some of the fitted parameters still differ substantially from their true values. At *N* = 1000, the fitted distribution becomes closer to the true stationary law, but differences in the recovered parameters remain. At *N* = 3000, the procedure selects the true value *a* = 4 and recovers 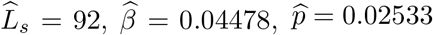, which are close to the ground truth values, while the fitted stationary distribution almost overlaps the true distribution.

We thus apply a standardized inference pipeline to experimental telomere length distributions with sample sizes *N* ≥ 1000 (Fig. 4).

**Figure 4.**
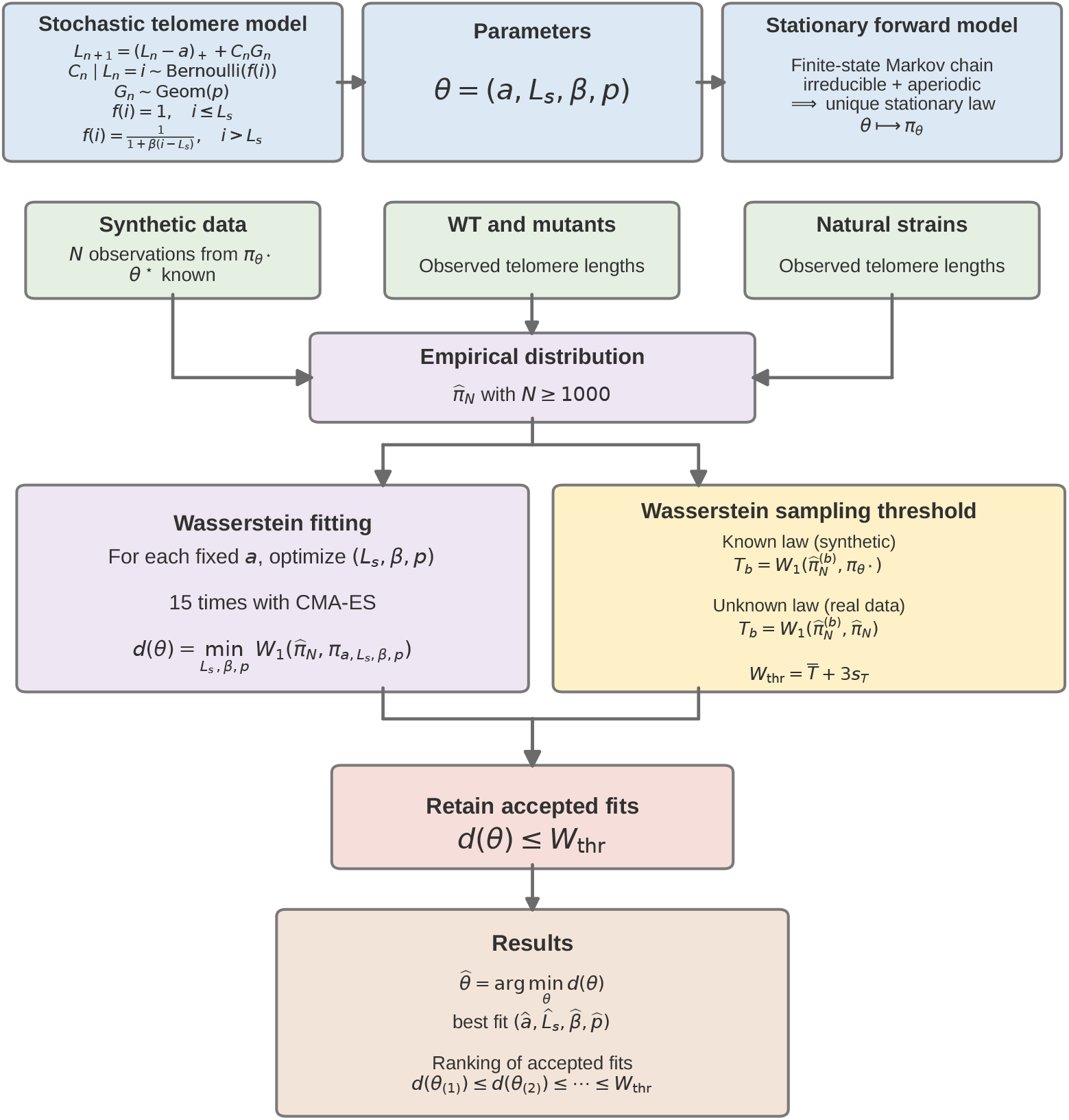
Scheme of the parameter inference pipeline. (Upper panel, blue boxes) The stochastic model is defined by four parameters and converges to a unique stationary distribution. (Green boxes) The datasets considered in this work comprise synthetic data used to assess the performance of the inference method; experimental data from wild-type and mutant strains affecting telomere length regulation [18], used to evaluate the biological relevance of the inferred parameters; and experimental data from natural yeast strains [19] displaying diverse telomere length distributions. The inference pipeline consists of optimizing the parameters (*L*_*s*_, *β, p*) using 15 independent CMA-ES runs, by minimizing the Wasserstein distance between the empirical distribution 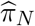 and the stationary model distribution *π*_*θ*_, for each fixed value of the shortening parameter *a*. A bootstrap-based threshold, 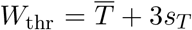, where 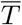 and *s*_*T*_ denote the mean and standard deviation of the bootstrap Wasserstein distances, respectively, is obtained using the known generating distribution for synthetic data and empirical resampling for experimental data. Parameter sets satisfying *d*(*θ*)≤*W*_thr_ are retained. The best-fitting parameter set and the accepted alternative solutions are then reported and ranked in increasing order of Wasserstein distance.

### The stationary distribution fits experimental telomere length distributions

We first apply our approach to telomere length distributions measured by ONT sequencing in the W303 laboratory strain and its *yku70*Δ and *rif1*Δ mutant derivatives [18]. Impairing the Ku complex leads to shorter telomeres [38–40], and, Rif1 being a negative regulator of telomere length, deleting *RIF1* results in longer telomeres [3, 41].

We found an excellent agreement between the experimental data and our best fits, for wild-type W303 and *yku70*Δ, and acceptable agreement for *rif1*Δ (Fig. 5A–C). In fact, for each genotype, several other fits passed the Wasserstein distance threshold and thus yielded acceptable sets of parameters. We therefore ranked the three best fits according to increasing Wasserstein distances, and present the corresponding parameters (Fig. 5A–C). We then repeated the parameter inference pipeline on independent biological replicates of all three genotypes [18]. We generally found good qualitative agreement for the wild-type replicates, although the best three fits did not quite cross the threshold W_thr_, due to a left tail in the experimental distribution, not accounted for in our model (Supp. Fig. S3A–E). For *yku70*Δ, the fit on the other replicates remained excellent (Supp. Fig. S3F and G). The fits of the other *rif1*Δ replicates were of variable quality, explained by an irregular shape of the experimental distributions (Supp. Fig. S3H and I). Nonetheless, the parameters found in these three genotypes are consistent with the known effects of *yku70*Δ and *rif1*Δ mutants on telomere homeostasis, and quantitative estimates from the literature (see Discussion).

**Figure 5.**
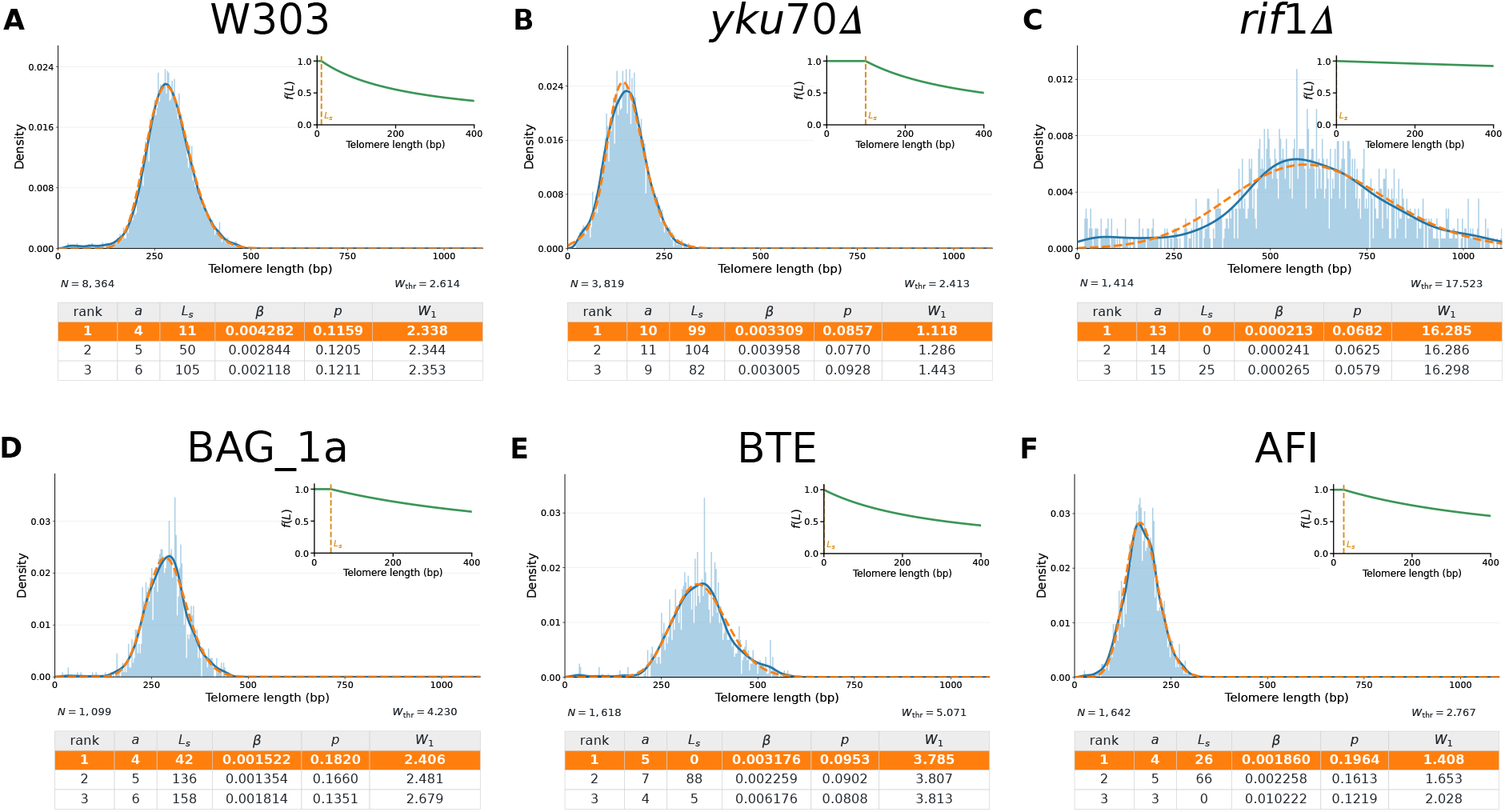
Estimation of parameters from experimental telomere length distributions. **(A–C)** Fits of telomere length distributions from wild-type laboratory strain W303 **(A)** and its mutant derivatives *yku70*Δ **(B)** and *rif1*Δ **(C). (D–F)** Fits of telomere length distributions from natural yeast isolates from diverse origins: BAG_1a sampled from tree bark in China **(D)**, BTE from an Italian bakery **(E)**, and AFI from French wine **(F)**. Blue histograms represent the experimental telomere length distributions, and the blue curves show the corresponding kernel density estimates. The superimposed orange curves represent the best-fitted stationary distributions. Insets display the corresponding telomerase recruitment function *f* (*L*), with the vertical dotted line indicating the estimated threshold *L*_*s*_. The tables below each panel report the parameter estimates (*a, L*_*s*_, *β, p*) and the Wasserstein distance *W*_1_ for the three highest-ranked fits, with the best fit highlighted in orange.

In *S. cerevisiae*, the steady-state distribution spans roughly 200–500 bp of telomere length in laboratory strains, but large variations across natural strains have been described [19, 42–45]. To gain insights into telomere length regulation across this diversity, we expanded our analyses to a panel of natural yeast strains, which were sequenced by ONT and which display diverse telomere length distributions, both in terms of average telomere length and overall shape [19, 45]. From the published dataset [19], we selected the 66 strains for which sufficient sample size was available (*N* ≥ 1000) and fitted our mathematical model (Fig. 5D-F and Supp. Fig. S4–S10). We show in (Fig. 5D-F) three natural yeast strains with diverse ecological origins and average telomere lengths to illustrate that our model captured the shape of these distributions, suggesting that the genetic diversity responsible for telomere length variations in all these strains (Fig. 5D-F and Supp. Fig. S4–S10) do not alter the fundamental architecture of telomere length regulation implemented in our model.

### Chromosome-end-specific telomere length distributions

Telomere length regulation ensures an overall equilibrium, but individual telomere lengths are highly variable across cells and across different extremities in a given cell population, not only in *S. cerevisiae*, but also in other eukaryotes, including humans [17, 19, 29, 46–48].

To investigate how telomeres across chromosome ends are differentially regulated, we applied our model to chromosome-end-specific telomere length distributions. We assigned each telomeric read in the dataset from Theulot et al. [18] to a chromosome end using subtelomeric sequence as anchor. In the wild-type strain, we observed robust differences in telomere length distributions between chromosome ends across replicates (Supp. Fig. S11). Telomere length distributions are less variable across chromosome ends in the *yku70*Δ mutant (Supp. Fig. S12), consistent with previous observations [19].

Because the sample size per chromosome end has to be large enough, we pooled together the independent replicates for wild-type and *yku70*Δ mutant from Theulot et al. [18]. The pooled data of the *rif1*Δ mutant did not contain enough telomeric reads for further analysis. The pooled data of the *yku70*Δ mutant contained an average sample size of *N* = 650.8 per chromosome end, a number which still allowed for reasonable fit accuracy despite not reaching 1000 (Fig. 3E–G).

In the wild-type, despite substantial differences between chromosome-end-specific distributions, the best-fit shortening rates remained remarkably close to the value found for the global distribution (Fig. 6A and Supp. Fig. S13–S16). Similarly, the best-fit values of p did not fluctuate much either, except for chromosome III left, indicating that the number of nucleotides added by telomeres would not be dependent of the specific chromosome end (Fig. 6D). In contrast, the best-fit values for *L*_*s*_ varied much more widely between 4 and 131, and while *β* values were less variable, the overall telomerase recruitment probability appeared strongly chromosome-end-specific (Fig. 6B and C). Interestingly, chromosome III left was associated with a particularly long average telomere length, as previously observed [17, 19, 45, 49–50]. The fitted parameters explained this longer telomere mostly by an increased probability of telomerase recruitment, driven by a high value of *L*_*s*_ (Fig. 6B and Supp. Fig. S13), consistent with recent findings showing increased Sir4–Ku–TLC1-dependent telomerase recruitment at this telomere due to the presence of the HML locus close to the telomere (Teplitz et al. 2026), but also by a greater elongation length (i.e. a small value of *p*).

**Figure 6.**
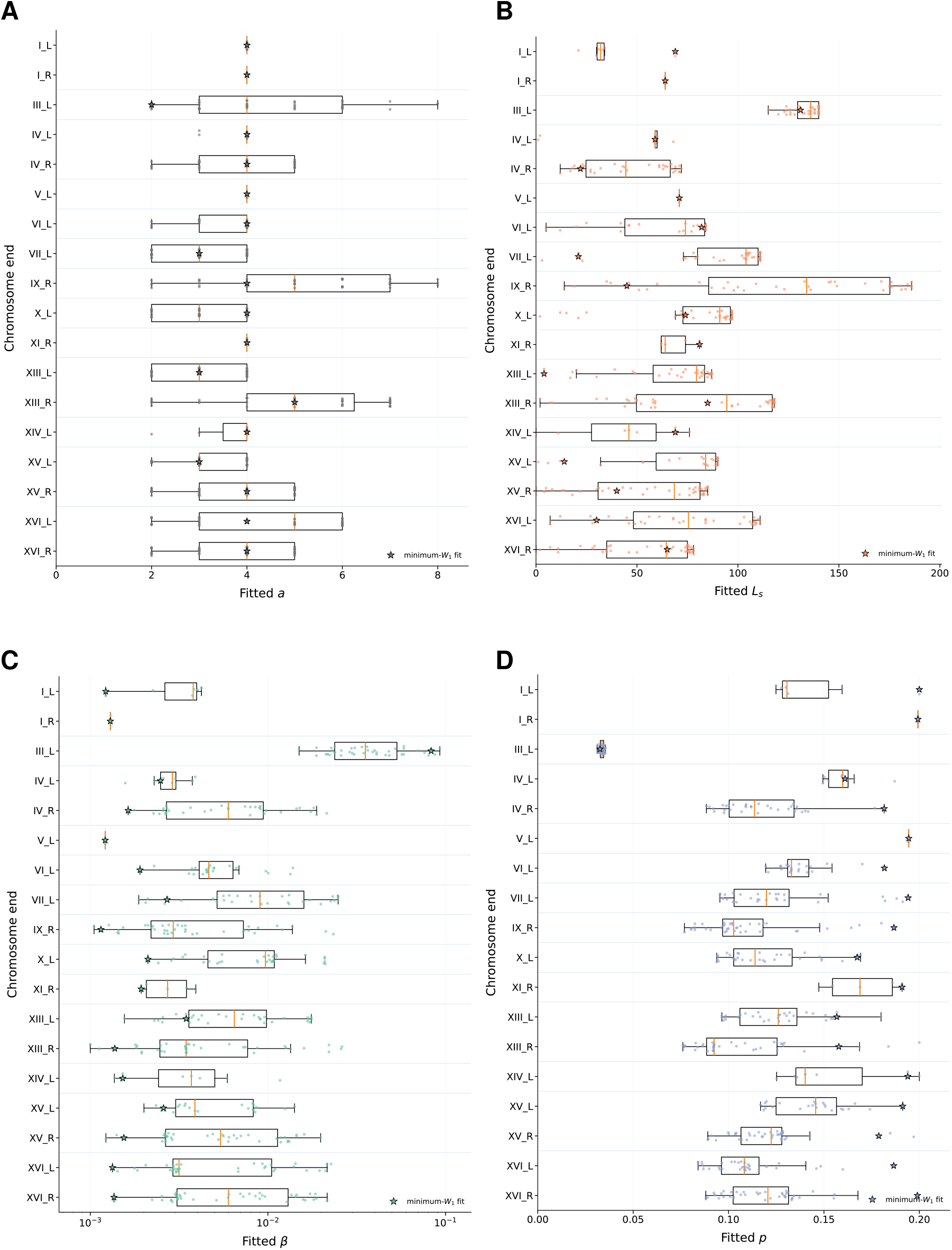
Chromosome-end-specific variability in fitted parameters for pooled W303 data. Measurements from six wild-type W303 replicates were pooled and analyzed separately for each of the 32 chromosome ends. CMA-ES optimization yielded acceptable fits for 18 chromosome ends. Boxplots show the distributions of fitted parameters across optimization runs: **(A)** *a*, **(B)** *L*_*s*_, **(C)** *β*, **(D)** *p*. Boxes indicate the interquartile range, central lines the median, and whiskers the spread of fitted values. The best-fit parameter is indicated by a star. The parameter *β* is shown on a logarithmic scale.

While the *yku70*Δ mutant displayed much less variability than wild-type (Supp. Fig. S11 and S12), the fitted parameters still varied across chromosome ends (Supp. Fig. S17). The best-fit shortening rates were similar to the value fitted with the global distribution, suggesting that the mechanism by which the Ku complex limits the 3′ overhang, and thus the shortening rate, acts similarly across all chromosome ends.

Overall, our pipeline can be used directly on chromosome-end-specific telomere length distributions and provide insights into how they might be differentially regulated.

## Discussion

In this work, we propose a mathematical modelling framework to investigate the molecular mechanisms of telomere dynamics using the steady-state telomere length distribution. Our approach is timely as long-read sequencing has recently enabled the measurement of telomeres at the single-molecule level and the analysis of the whole telomere length distribution at an unprecedented resolution [17, 19, 51]. The inverse problem of inference biochemical parameters from this distribution is not trivial, however, and we first laid solid mathematical foundations showing the convergence of the Markov chain representing telomere dynamics towards a steady-state distribution with an explicit expression, and the theoretical identifiability of the inference procedure.

From a numerical perspective and with synthetic data, we found that minimizing the Wasserstein distance led to optimal parameters increasingly close to their true values, provided a large enough sample size. While the best-fit parameters agreed very well with the ground truth, other acceptable fits for *a* and *p* were also equal or quite close to the true values. However, other acceptable fits for *L*_*s*_ and *β* were more variable, indicating that these parameters might be more difficult to retrieve or might compensate for each other in practice. Therefore, for sample sizes that are limiting, the confidence on the estimation of these two parameters might be lower, which will be reflected in the ranked solutions we provide in our pipeline.

The application of our approach to experimental data confirmed its validity. For the wild-type yeast strain from [18], the shortening rate of the best fit was found to be 4, consistent with estimated shortening rates in the literature [8, 52–55]. The best-fit value of *p* = 0.1958 corresponded to an average elongation of approximately 5 nucleotides per event, which is smaller than the average of 30–40 bp measured experimentally [14]. This discrepancy can be explained by two aspects of our implementation of telomere elongation. First, telomerase has been shown to be processive *in vivo*, that is, able to add long telomeric sequences in a single event through dissociation and reassociation with the telomere, but mostly at telomeres shorter than 125 bp [56]. We did not include this property in our model for simplicity and because most telomeres in a wild-type strain are longer than 125 bp. Excluding these short telomeres in the experimental data from Teixeira et al. [14] would reduce the discrepancy we observed. Second, for elongation, we used a geometric distribution with constant probability *p*, which allows for all elongation lengths of at least one nucleotide. *In vivo*, however, after annealing of the TLC1 RNA template to the 3′ overhang, telomerase tends to synthesize the central part of the template without interruption, leading to the frequent incorporation of at least 6–7 nucleotides and up to 15 nucleotides [57], and therefore to very rare incorporation of five or less nucleotides. Implementing such a minimum number of incorporated nucleotides would mechanically result in a longer average elongation, with compensatory changes in *f* (*L*) to decrease the frequency of elongation. As a consequence of this compensation, the best fit values of *L*_*s*_ and *β* did not correspond well with experimental data measuring the probability of single telomerase extension events as a function of telomere length [14]. We must also note that the experimental data for *f* (*L*) showed much more inherent variability, making it difficult to assess the biological relevance of our fits. We conclude that, although the exact fitted values of *p, L*_*s*_, and *β* might not be directly interpretable biologically, their relative differences across strains and mutants, and across chromosome ends should be relevant biologically.

Telomeres in Ku mutants displays longer 3′ overhangs, which would lead to greater telomere shortening rate [58–61]. In addition, the Ku complex also contributes to telomerase recruitment through its interaction with Rap1-bound Sir4 and TLC1 ([39–40, 62–63]). The shorter telomeres in a mutant of the Ku complex thus stem from two additive effects: greater shortening rate and decreased telomerase recruitment. Our best-fit parameters recapitulate these effects with *a* = 8 and a *f* (*L*_*s*_, *β*) curve displaying decreased telomerase recruitment probability overall compared to the wild-type strain. Rif1 contributes the length-dependent negative regulation of telomerase recruitment [41, 55]. Consistently, the best-fit parameters yielded *f* (*L*_*s*_, *β*) corresponding to an increased probability of telomerase recruitment at any telomere length compared to wild-type, in agreement with experimental data measuring telomerase elongation events in *rif1*Δ [14]. However, our model failed to fit the experimental distribution of two replicates out of three of *rif1*Δ and we suspect that telomere length regulation is drastically altered in this mutant, invalidating the application of our model.

The ability to access chromosome-end-specific telomere length distributions using long-read sequencing opens new research avenues into the cis or local regulators of telomere length [17, 19, 46, 50]. We found that in the wild-type W303 strain, most of the differences across chromosome ends stems from how telomerase is recruited to the telomere and not from the shortening rate or telomerase elongation. These differences are significantly dampened in the *yku70*Δ mutant, suggesting that the Ku-dependent telomerase recruitment pathway, possibly with variable activity, might be a major contributor of the inter-extremity differences, as it was demonstrated for the long chromosome III L telomere [50]. Our pipeline thus provides insights into the parameters that are potentially variable across chromosome ends, thus generating hypotheses to be further tested experimentally.

To conclude, our work demonstrates that mathematical modelling of telomere length dynamics is a powerful approach to relate the biochemical parameters controlling individual telomere evolution over cell divisions to the steady-state length distribution, thus advancing our understanding of telomere regulation. In addition, with minor adaptations, our approach should be applicable to telomere length distributions from other species as well, including humans.

## Material and Methods

### Telomere length datasets and preprocessing

Telomere lengths were obtained from long-read sequencing data [18–19]. Identification and extraction of telomeric reads, together with the upstream sequence-processing procedure, followed the telomere analysis pipeline described in [19]. The analyses presented here therefore started from the resulting telomere length measurements and focused on the mathematical and statistical inference of telomere length regulation parameters.

Before model fitting, outliers corresponding to unusually large telomere lengths were removed using an interquartile-range criterion. Let *Q*_1_ and *Q*_3_ denote the first and third quartiles, respectively, and let IQR = *Q*_3_ − *Q*_1_. An observation *L* was excluded when *L > Q*_3_ + 2 IQR. All sample sizes, empirical distributions, parameter estimation procedures, and bootstrap calculations described below were based on the resulting filtered observations. For chromosome extremity analyses, groups containing fewer than 1000 observations were excluded from model fitting.

### Stationary distribution and Wasserstein objective

For a parameter vector *θ* = (*a, β, L*_*s*_, *p*), the telomere model defines a discrete-state Markov chain whose stationary distribution, denoted by *π*_*θ*_, represents the predicted steady state telomere length distribution. Rather than simulating individual telomere trajectories during optimization, the stationary distribution was calculated numerically for each candidate parameter vector and directly compared with the experimental distribution.

Numerical calculations were performed on the truncated state space *S* = {0, 1, …, *L*_max_}. The value of *L*_max_ was chosen above the relevant range of the experimental telomere-length distributions so that truncation had negligible influence on the inferred distribution.

For each parameter set, the corresponding stationary telomere length distribution was computed numerically from the Markov model. The numerical procedure and the analysis of convergence toward stationarity are described in detail in the Supp. Text.

Let *π*_exp_ denote the empirical distribution obtained from the observed telomere lengths. Model fitting was based on the one dimensional Wasserstein distance *W*_1_(*π*_exp_, *π*_*θ*_).

The Wasserstein distance was used because it compares the complete shape and position of the two telomere length distributions while explicitly accounting for differences in telomere length. In particular, moving probability mass between nearby lengths incurs a smaller discrepancy than moving it between distant lengths, making it well suited to comparing empirical and modelled telomere length distributions.

### Parameter estimation using CMA-ES

Parameter estimation was performed by minimizing *W*_1_(*π*_exp_, *π*_*θ*_) using the Covariance Matrix Adaptation Evolution Strategy (CMA-ES) [37]. CMA-ES is a stochastic derivative-free optimization method developed for nonlinear and non-convex continuous optimization problems. At each generation, the algorithm samples a population of candidate solutions from an adaptive multivariate Gaussian distribution, evaluates and ranks these candidates according to the objective function, and updates the search distribution using the best performing solutions. Adaptation of the covariance matrix allows the search to progressively account for differences in parameter scales and correlations between parameters [37].

CMA-ES was appropriate for the present problem because each evaluation of the objective function requires the numerical computation of a stationary distribution and no simple analytical expression for the gradient of the Wasserstein objective is available. In addition, the parameter landscape may contain flat regions and compensatory relationships between parameters, making a derivative free global optimization strategy preferable to a single local gradient based optimization.

Because *a* is an integer-valued parameter and the identifiability analysis showed that inference of the remaining parameters must be considered conditionally on *a*, it was treated by profiling rather than as a continuous CMA-ES coordinate. We considered *a* ∈ {2, …, 20}. For each fixed value of *a*, CMA-ES was used to optimize the remaining parameter vector (*β, L*_*s*_, *p*). For the experimental data analyses, the general parameter search domain was 10^*−*4^ ≤ *β* ≤ 0.20, 0.003 ≤ *p* ≤ 0.20, with 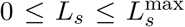. The upper bound of *L*_*s*_ was constrained using the lower tail of the empirical telomere-length distribution, 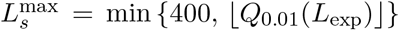, where *Q*_0.01_(*L*_exp_) denotes the empirical first percentile of the corresponding dataset. This restriction prevents *L*_*s*_ from exploring regions poorly supported by the experimentally observed short telomere distribution.

For each fixed value of *a*, 15 independent CMA-ES optimizations were performed using different initializations. Since fixing *a* leaves three continuous parameters to estimate, (*β, L*_*s*_, *p*), the standard CMA-ES population-size rule, *λ* = 4 + ⌊3 ln(*d*)⌋, gives *λ* = 7 for *d* = 3 [37]. This value is a default recommendation rather than a strict requirement, and moderately increasing the population size can improve exploration of the parameter space and reduce sensitivity to local optima, at the cost of additional function evaluations [37]. We therefore used a slightly larger population size of *λ* = 12 candidate parameter vectors per generation, which provides greater search diversity while remaining computationally moderate for this optimization problem. Each optimization was run for at most 100 generations. Consequently, a single CMA-ES run involved at most approximately 12 × 100 = 1200 objective-function evaluations, unless convergence occurred earlier. Multiple independent runs were used to further reduce sensitivity to stochastic initialization and to explore alternative parameter combinations associated with comparable objective values.

For each fixed *a*, the solution attaining the smallest Wasserstein distance among the independent runs was taken as the profile optimum for that value of *a*. The overall minimum was then obtained by comparing the profile optima across all tested values of *a*. The remaining optimization runs were retained to evaluate the variability of parameter estimates and to investigate the existence of alternative parameter combinations yielding similarly close stationary distributions.

### Bootstrap based Wasserstein threshold

A small Wasserstein distance does not by itself determine whether the residual discrepancy between an estimated stationary distribution and the experimental distribution is larger than could reasonably arise from finite sampling. We therefore constructed an empirical bootstrap reference distribution independently for each dataset.

For a dataset containing *n* observed telomere lengths after preprocessing, we generated *B* = 1000 bootstrap samples, each containing *n* observations drawn with replacement from the original dataset. For bootstrap replicate *b* = 1, …, *B*, we constructed the corresponding empirical distribution 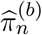 and computed 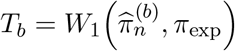. The collection *T*_1_, …, *T*_*B*_ provides an empirical estimate of the magnitude of Wasserstein variability expected from finite sampling at the observed sample size and for the specific shape of the corresponding experimental distribution.

The empirical mean of the bootstrap distances was calculated as 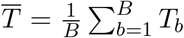, and their standard deviation as 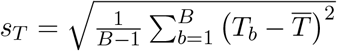. The bootstrap-based Wasserstein threshold was defined as 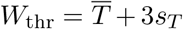, where 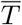 and *s*_*T*_ represent the empirical mean and sample standard deviation of the bootstrap Wasserstein distances, respectively.

This procedure was applied independently to each analysis unit, using its own empirical observations and sample size. Thus, replicate level, pooled strain, chromosome level, chromosome extremity, and natural strain distributions each received their own bootstrap reference threshold. This was particularly important for chromosome extremity analyses because sample sizes can differ substantially among chromosome ends.

For an estimated parameter vector 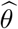, the model discrepancy 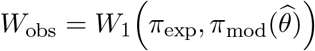 was compared with *W*_thr_. Fits satisfying *W*_obs_ ≤ *W*_thr_ were considered compatible with the magnitude of variability expected from finite sampling, whereas larger distances indicated a model data discrepancy exceeding this empirical reference scale.

### Validation using synthetic data

The complete inference procedure was first evaluated on synthetic datasets for which the generating parameters were known. For a prescribed parameter vector 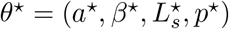, the corresponding stationary distribution 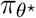 was computed numerically. Independent samples of different sizes *N* were then generated from this stationary distribution and treated as empirical datasets.

The same inference procedure used for the experimental data was applied to these synthetic samples without providing the generating parameters to the optimization algorithm. In particular, inference included the fixed-*a* scan, repeated CMA-ES optimizations, numerical calculation of the stationary distribution, and Wasserstein-based comparison between observed and predicted distributions.

### Computational implementation and high-performance computing

The numerical inference pipeline was implemented in Python 3. Numerical and tabular calculations used primarily `NumPy` and `pandas`; statistical calculations, including the one dimensional Wasserstein distance, used `SciPy`; CMA-ES optimization was performed with the Python `cma` package (*pycma*); and graphical analyses were produced using `Matplotlib`. Jupyter notebooks were used for exploratory analyses, statistical summaries, quality control, and generation of publication figures, while computationally intensive model fitting procedures were implemented in Python.

The large scale parameter estimation calculations were performed on the IFB Core Cluster of the Institut Français de Bioinformatique (IFB) [64] using the SLURM workload manager. The cluster was used in particular for the repeated CMA-ES optimizations required for the fixed *a* profiles and for the systematic analyses of independent strains, biological replicates, pooled datasets, chromosomes, and chromosome extremities.

Optimization results were stored in structured output tables containing the fitted parameters, Wasserstein distances, sample sizes, optimization identifiers, and associated fitting information. Stationary distributions and parameter profiles required for downstream analyses were stored separately. This separation allowed computationally expensive parameter estimation to be performed on the cluster while subsequent filtering, statistical summaries, bootstrap comparisons, and graphical analyses could be conducted without rerunning the optimization procedure.

The same stationary-distribution solver, Wasserstein criterion, optimization strategy, and parameter definitions were used for both synthetic and experimental datasets to ensure direct comparability between validation and biological applications.

## Data availability

The experimental telomere length data analyzed in this study were obtained from previously published Oxford Nanopore sequencing datasets. Data from 100 natural *Saccharomyces cerevisiae* strains were obtained from [19], with sequencing data available in the European Nucleotide Archive (ENA) under accession number PRJEB89249 (https://www.ebi.ac.uk/ena/browser/view/PRJEB89249).

Sequencing data for laboratory wild-type and mutant strains were obtained from Theulot et al. [18], available in the ENA under accession number PRJEB76824 (https://www.ebi.ac.uk/ena/browser/view/PRJEB76824).

The Python code implementing the mathematical model, stationary distribution computations, parameter estimation, bootstrap procedures, and figure generation, together with the processed datasets required to reproduce the analyses, will be made available on GitHub at https://github.com/kvgg7/TL_SC.

## Acknowledgement

Research in ZX’s laboratory was supported by Agence Nationale de la Recherche (ANR-24-CE12-7740-01 and ANR-24-CE12-0083-03). The PhD fellowship of VGG is supported by Mission pour les Initiatives Transverses et Interdisciplinaires (MITI) of Centre National de la Recherche Scientifique (CNRS). VGG and MD acknowledge the support of the Chaire MMB (Modélisation Mathématique et Biodiversité). This work was also supported by the France 2030 plan operated by Agence Nationale de la Recherche (ANR-23-EXMA-0005).

The authors warmly thank Chloé Audebert for stimulating discussions and her continued interest in this research, as well as Clotilde Garrido for her help in sequencing data processing. VGG also thanks Oskar Girardin for a helpful discussion on the CMA-ES optimization algorithm.

## Author contributions

VGG performed methodology, investigation, formal analysis, and writing the original draft. MD performed investigation, conceptualization, supervision and writing the original draft. ZX performed conceptualization, supervision, and writing the original draft. All authors reviewed and edited the manuscript.

## References

[1] R. J. Wellinger and V. A. Zakian. Everything you ever wanted to know about saccharomyces cerevisiae telomeres: beginning to end. Genetics, 191:1073–1105, 2012.

[2] T. de Lange. Shelterin-mediated telomere protection. Annu Rev Genet, 52:223–247, 2018.

[3] S. Marcand, E. Gilson, and D. Shore. A protein-counting mechanism for telomere length regulation in yeast. Science, 275:986–990, 1997.

[4] D. L. Levy and E. H. Blackburn. Counting of rif1p and rif2p on saccharomyces cerevisiae telomeres regulates telomere length. Mol Cell Biol, 24:10857–10867, 2004.

[5] J. D. Watson. Origin of concatemeric t7 dna. Nat New Biol, 239:197–201, 1972.

[6] A. M. Olovnikov. A theory of marginotomy. The incomplete copying of template margin in enzymic synthesis of polynucleotides and biological significance of the phenomenon. J Theor Biol, 41:181–190, 1973.

[7] J. Lingner, J. P. Cooper, and T. R. Cech. Telomerase and dna end replication: no longer a lagging strand problem? science 269: 1533–1534. 1995.

[8] V. Lundblad and J. W. Szostak. A mutant with a defect in telomere elongation leads to senescence in yeast. Cell, 57:633–643, 1989.

[9] S. Enomoto, L. Glowczewski, and J. Berman. Mec3, mec1, and ddc2 are essential components of a telomere checkpoint pathway required for cell cycle arrest during senescence in saccharomyces cerevisiae. Mol Biol Cell, 13:2626–2638, 2002.

[10] A. S. Ijpma and C. W. Greider. Short telomeres induce a dna damage response in saccharomyces cerevisiae. Mol Biol Cell, 14:987–1001, 2003.

[11] C. W. Greider and E. H. Blackburn. Identification of a specific telomere terminal transferase activity in tetrahymena extracts. Cell, 43:405–413, 1985.

[12] M. M. Ouellette, M. Liao, B. S. Herbert, M. Johnson, S. E. Holt, H. S. Liss, J. W. Shay, and W. E. Wright. Subsenescent telomere lengths in fibroblasts immortalized by limiting amounts of telomerase. J Biol Chem, 275:10072–10076, 2000.

[13] M. T. Hemann, M. A. Strong, L. Y. Hao, and C. W. Greider. The shortest telomere, not average telomere length, is critical for cell viability and chromosome stability. Cell, 107:67–77, 2001.

[14] M. T. Teixeira, M. Arneric, P. Sperisen, and J. Lingner. Telomere length homeostasis is achieved via a switch between telomerase-extendible and -nonextendible states. Cell, 117: 323–335, 2004.

[15] B. Britt-Compton, R. Capper, J. Rowson, and D. M. Baird. Short telomeres are preferentially elongated by telomerase in human cells. FEBS Lett, 583:3076–3080, 2009.

[16] Z. Xu, K. Dao Duc, D. Holcman, and M. T. Teixeira. The length of the shortest telomere as the major determinant of the onset of replicative senescence. Genetics, 194:847–857, 2013.

[17] S. L. Sholes, K. Karimian, A. Gershman, T. J. Kelly, W. Timp, and C. W. Greider. Chromosome-specific telomere lengths and the minimal functional telomere revealed by nanopore sequencing. Genome Res, 32:616–628, 2022.

[18] B. Theulot, A. Tourancheau, E. Simonin Chavignier, E. Jean, J. M. Arbona, B. Audit, O. Hyrien, L. Lacroix, and B. Le Tallec. Telomere-to-telomere dna replication timing profiling using single-molecule sequencing with nanotiming. Nat Commun, 16:242, 2025.

[19] C. Garrido, C. Gomez-Munoz, E. Kornobis, N. Agier, O. Ilioaia, G. Fischer, and Z. Xu. Natural diversity of telomere length distributions across 100 saccharomyces cerevisiae strains. Genome Res, 2026. doi: 10.1101/gr.281132.125.

[20] K. B. Blagoev. Cell proliferation in the presence of telomerase. PLoS ONE, 4:e4622, 2009.

[21] A. Kowald. Possible mechanisms for the regulation of telomere length. Journal of Molecular Biology, 273:814–825, 1997.

[22] K. Oexle. Telomere length distribution and southern blot analysis. J Theor Biol, 190:369–377, 1998.

[23] I. A. Rodriguez-Brenes and C. S. Peskin. Quantitative theory of telomere length regulation and cellular senescence. Proc Natl Acad Sci U S A, 107:5387–5392, 2010.

[24] K. D. Duc and D. Holcman. Computing the length of the shortest telomere in the nucleus. Phys Rev Lett, 111:228104, 2013.

[25] B. V. Hirt, J. A. Wattis, and S. P. Preston. Modelling the regulation of telomere length: the effects of telomerase and g-quadruplex stabilising drugs. J Math Biol, 68:1521–1552, 2014.

[26] S. Eugene, T. Bourgeron, and Z. Xu. Effects of initial telomere length distribution on senescence onset and heterogeneity. J Theor Biol, 413:58–65, 2017.

[27] J. Z. Farkas and G. F. Webb. Mathematical analysis of a clonal evolution model of tumour cell proliferation. Journal of Evolution Equations, 17:275–308, 2017.

[28] K. H. Lee and M. Kimmel. Stationary distribution of telomere lengths in cells with telomere length maintenance and its parametric inference. Bull Math Biol, 82:150, 2020.

[29] J. Shampay and E. H. Blackburn. Generation of telomere-length heterogeneity in saccharomyces cerevisiae. Proc Natl Acad Sci U S A, 85:534–538, 1988.

[30] J. R. Norris. Markov Chains. Cambridge University Press, 1997.

[31] M. Benaïm and N. El Karoui. Promenade aléatoire: chaînes de Markov et simulations, martingales et stratégies. Editions de l’Ecole Polytechnique, 2005. ISBN 9782730211680.

[32] Cédric Villani. Optimal Transport: Old and New, volume 338 of Grundlehren der mathematischen Wissenschaften. Springer, 2009. doi: 10.1007/978-3-540-71050-9.

[33] Filippo Santambrogio. Optimal Transport for Applied Mathematicians: Calculus of Variations, PDEs, and Modeling, volume 87 of Progress in Nonlinear Differential Equations and Their Applications. Birkhäuser / Springer, Cham, 2015. ISBN 9783319208275. doi: 10.1007/978-3-319-20828-2.

[34] L. Ljung and T. Glad. On global identifiability for arbitrary model parametrizations. Automatica, 30(2):265–276, 1994. doi: 10.1016/0005-1098(94)90029-9.

[35] F.-G. Wieland, A. L. Hauber, M. Rosenblatt, C. Tönsing, and J. Timmer. On structural and practical identifiability. Current Opinion in Systems Biology, 25:60–69, 2021. doi: 10.1016/j.coisb.2021.03.005.

[36] M. J. Simpson and O. J. Maclaren. Profile-wise analysis: A profile likelihood-based workflow for identifiability analysis, estimation, and prediction with mechanistic mathematical models. PLoS Computational Biology, 19(9):e1011515, 2023. doi: 10.1371/journal.pcbi.1011515.

[37] N. Hansen. The cma evolution strategy: A tutorial, 2023.

[38] S. E. Porter, P. W. Greenwell, K. B. Ritchie, and T. D. Petes. The dna-binding protein hdf1p (a putative ku homologue) is required for maintaining normal telomere length in saccharomyces cerevisiae. Nucleic Acids Res, 24:582–585, 1996.

[39] A. E. Stellwagen, Z. W. Haimberger, J. R. Veatch, and D. E. Gottschling. Ku interacts with telomerase rna to promote telomere addition at native and broken chromosome ends. Genes Dev, 17:2384–2395, 2003.

[40] T. S. Fisher, A. K. Taggart, and V. A. Zakian. Cell cycle-dependent regulation of yeast telomerase by ku. Nat Struct Mol Biol, 11:1198–1205, 2004.

[41] C. F. Hardy, L. Sussel, and D. Shore. A rap1-interacting protein involved in transcriptional silencing and telomere length regulation. Genes Dev, 6:801–814, 1992.

[42] R. M. Walmsley and T. D. Petes. Genetic control of chromosome length in yeast. Proc Natl Acad Sci U S A, 82:506–510, 1985.

[43] G. Liti, S. Haricharan, F. A. Cubillos, A. L. Tierney, S. Sharp, A. A. Bertuch, L. Parts, E. Bailes, and E. J. Louis. Segregating yku80 and tlc1 alleles underlying natural variation in telomere properties in wild yeast. PLoS Genet, 5:e1000659, 2009.

[44] M. D’Angiolo, J. X. Yue, M. De Chiara, B. P. Barre, M. J. Giraud Panis, E. Gilson, and G. Liti. Telomeres are shorter in wild saccharomyces cerevisiae isolates than in domesticated ones. Genetics, 223, 2023.

[45] S. O’Donnell, J. X. Yue, O. A. Saada, N. Agier, C. Caradec, T. Cokelaer, M. De Chiara, S. Delmas, F. Dutreux, T. Fournier, et al. Telomere-to-telomere assemblies of 142 strains characterize the genome structural landscape in saccharomyces cerevisiae. Nat Genet, 55: 1390–1399, 2023.

[46] K. Karimian, A. Groot, V. Huso, R. Kahidi, K. T. Tan, S. Sholes, R. Keener, J. F. McDyer, J. K. Alder, H. Li, et al. Human telomere length is chromosome end-specific and conserved across individuals. Science, 384:533–539, 2024.

[47] G. M. Teplitz, E. Pasquier, E. Bonnell, E. De Laurentiis, L. Bartle, J. F. Lucier, S. Sholes, C. W. Greider, and R. J. Wellinger. A mechanism for telomere-specific telomere length regulation. bioRxiv, 2024. doi: 10.1101/2024.06.12.598646.

[48] I. Biocanin, A. Edera, C. Garrido, A. Vidal, D. Dardalhon-Cuménal, R. Burns, K. Fritschi, F. A. Rabanal, J. Fajkus, C. Bourbousse, et al. Centromeric origin of arabidopsis interstitial telomeric repeat domains. bioRxiv, 2026.

[49] L. L. Button and C. R. Astell. The saccharomyces cerevisiae chromosome iii left telomere has a type x, but not a type y’, ars region. Mol Cell Biol, 6:1352–1356, 1986.

[50] G. M. Teplitz, E. Pasquier, E. Bonnell, E. De Laurentiis, L. Bartle, J. F. Lucier, G. St-Laurent, D. S. Dawson, and R. J. Wellinger. A mechanism for telomere-specific telomere length regulation. Nucleic Acids Res, 54, 2026.

[51] L. Dudragne, C. Garrido, O. Ilioaia, J. S. Bernardes, and Z. Xu. Transient telomere uncapping triggers telomeric and subtelomeric rearrangements. EMBO Reports, 2026. doi: 10.1038/s44319-026-00717-4.

[52] M. S. Singer and D. E. Gottschling. Tlc1: template rna component of saccharomyces cerevisiae telomerase. Science, 266:404–409, 1994.

[53] T. S. Lendvay, D. K. Morris, J. Sah, B. Balasubramanian, and V. Lundblad. Senescence mutants of saccharomyces cerevisiae with a defect in telomere replication identify three additional est genes. Genetics, 144:1399–1412, 1996.

[54] C. I. Nugent, T. R. Hughes, N. F. Lue, and V. Lundblad. Cdc13p: a single-strand telomeric dna-binding protein with a dual role in yeast telomere maintenance. Science, 274:249–252, 1996.

[55] S. Marcand, V. Brevet, and E. Gilson. Progressive cis-inhibition of telomerase upon telomere elongation. EMBO J, 18:3509–3519, 1999.

[56] M. Chang, M. Arneric, and J. Lingner. Telomerase repeat addition processivity is increased at critically short telomeres in a Tel1-dependent manner in Saccharomyces cerevisiae. Genes & Development, 21:2485–2494, 2007.

[57] K. Forstemann and J. Lingner. Molecular basis for telomere repeat divergence in budding yeast. Molecular and Cellular Biology, 21:7277–7286, 2001.

[58] S. Gravel, M. Larrivee, P. Labrecque, and R. J. Wellinger. Yeast ku as a regulator of chromosomal dna end structure. Science, 280:741–744, 1998.

[59] R. M. Polotnianka, J. Li, and A. J. Lustig. The yeast ku heterodimer is essential for protection of the telomere against nucleolytic and recombinational activities. Curr Biol, 8:831–834, 1998.

[60] D. Bonetti, M. Clerici, S. Anbalagan, M. Martina, G. Lucchini, and M. P. Longhese. Shelterinlike proteins and yku inhibit nucleolytic processing of saccharomyces cerevisiae telomeres. PLoS Genet, 6:e1000966, 2010.

[61] J. Soudet, P. Jolivet, and M. T. Teixeira. Elucidation of the dna end-replication problem in saccharomyces cerevisiae. Molecular Cell, 53:954–964, 2014.

[62] S. E. Peterson, A. E. Stellwagen, S. J. Diede, M. S. Singer, Z. W. Haimberger, C. O. Johnson, M. Tzoneva, and D. E. Gottschling. The function of a stem-loop in telomerase rna is linked to the dna repair protein ku. Nat Genet, 27:64–67, 2001.

[63] H. Chen, J. Xue, D. Churikov, E. P. Hass, S. Shi, L. D. Lemon, P. Luciano, A. A. Bertuch, D. C. Zappulla, V. Geli, et al. Structural insights into yeast telomerase recruitment to telomeres. Cell, 172:331–343 e313, 2018.

[64] Institut Français de Bioinformatique. IFB Core Cluster. https://doc.cluster.france-bioinformatique.fr/, 2026. Institut Français de Bioinformatique (IFB), ELIXIR France.

