## Supplementary Information for "Telomere dynamics parameters inferred from mathematical modeling of telomere length distribution"

### Part I. Supplementary Text

#### S1 Stochastic description of telomere length evolution

##### S1.1 Update equation and parameter definitions

As stated in Eq. (1) of the main article, the model describes the telomere length $L_n \in \{0, 1, 2, \ldots\}$ at generation $n$. Its general update rule is

$$L_{n+1} = (L_n - S_n)_+ + C_n Y_n, \tag{S1.1}$$

where $(x)_+ = \max\{x, 0\}$. For fitting, the shortening amplitude is assumed constant, $S_n = a$, with integer $a \geq 1$. The recruitment indicator $C_n$ is Bernoulli conditional on the current telomere length; whenever recruitment occurs, the positive elongation length $Y_n$ follows a geometric distribution. More precisely,

$$\mathbb{P}(C_n = 1 \mid L_n = i) = f(i), \tag{S1.2}$$

$$\mathbb{P}(Y_n = k) = p(1-p)^{k-1}, \qquad k = 1, 2, \ldots, \quad 0 < p < 1. \tag{S1.3}$$

Conditionally on $L_n$, the recruitment and elongation variables are independent, and the same update mechanism is used at every generation. The update rule is the one written in Eq. (3). The fitted recruitment function is

$$f(i) = \begin{cases} 1, & i \leq L_s, \\ \dfrac{1}{1 + \beta(i - L_s)}, & i > L_s, \end{cases} \qquad L_s \in \{0, 1, 2, \ldots\}, \quad \beta > 0. \tag{S1.4}$$

Throughout this supplement, the parameter vector is ordered as in the main model definition:

$$\theta = (a, L_s, \beta, p). \tag{S1.5}$$

The four coordinates respectively represent shortening per generation, the recruitment plateau endpoint, the decline in recruitment above the plateau, and the geometric elongation parameter.The expression $\frac{1}{p}$ is the model's mean extension length *conditional on recruitment.*

##### S1.2 Markov property and transition probabilities

Given $L_n = i$, the next state is obtained by first replacing $i$ with $r_i = (i-a)_+$ and then either leaving it unchanged (no recruitment) or adding $Y_n$ (recruitment). As the conditional distribution of the next state depends on the past only through $i$, $(L_n)_{n\geq 0}$ is a time-homogeneous Markov chain. Its transition kernel is

$$\boxed{\begin{aligned} P_\theta(i,j) &= (1-f(i))\mathbf{1}_{\{j=r_i\}} \\ &\quad + f(i)\,p(1-p)^{j-r_i-1}\mathbf{1}_{\{j\geq r_i+1\}}. \end{aligned}} \tag{S1.6}$$

The kernel is stochastic because $\sum_{j\geq 0} P_\theta(i,j) = (1-f(i)) + f(i)\sum_{k\geq 1} p(1-p)^{k-1} = 1$.

##### S1.3 Conditional moments and qualitative parameter compensation

The conditional expected change is

$$\mathbb{E}[L_{n+1} - L_n \mid L_n = i] = (i-a)_+ - i + \frac{f(i)}{p}. \tag{S1.7}$$

In particular, for $i \geq a$,

$$\mathbb{E}[L_{n+1} - L_n \mid L_n = i] = -a + \frac{f(i)}{p}. \tag{S1.8}$$

Furthermore, using $\mathbb{E}[Y_n^2] = (2-p)/p^2$,

$$\operatorname{Var}(C_n Y_n \mid L_n = i) = \frac{f(i)(2-p-f(i))}{p^2}. \tag{S1.9}$$

Thus different parameters can partially compensate *in the local mean drift*, without making the full transition laws equal. This explains why good fits to a finite stationary sample need not produce precise estimates of each parameter; it is not a proof of non-identifiability.

#### S2 Existence and convergence of the stationary distribution

##### S2.1 Existence of one positive recurrent, aperiodic class

Because $f(i) = [1+\beta(i-L_s)]^{-1}$ for $i > L_s$, one has $f(i) \to 0$ as $i \to \infty$. Consequently Eq. (S1.7) implies that, for sufficiently large $i$,

$$\mathbb{E}[L_{n+1} - L_n \mid L_n = i] \leq -a/2 < 0. \tag{S2.1}$$

This negative drift prevents the process from maintaining arbitrarily large lengths indefinitely without returning toward a finite range.

**Proposition S2.1** (Existence, uniqueness, and convergence)**.** *For every $a > 1$, finite integer $L_s \geq 0$, $\beta > 0$, and $0 < p < 1$, the markov chain defined by Eq.* (S1.6) *has a unique stationary probability distribution $\pi_\theta$. The chain has exactly one closed communicating class, and that class is positive recurrent and aperiodic. From every initial state, the distribution of $L_n$ converges to $\pi_\theta$.*

*Proof.* **Step 1: there is at most one closed communicating class.** Choose an integer $K > \max\{L_s, a\} + a$. From any $i \leq K + a - 1$, we have $r_i \leq K - 1$. Therefore a recruitment event followed by elongation by exactly $K - r_i \geq 1$ bp moves the chain to state $K$ in one step, with positive probability

$$f(i)p(1-p)^{K-r_i-1} > 0.$$

From any $i \geq K + a$, one may first have consecutive non-recruitment events. Every intermediate length remains greater than $L_s$ until the chain enters $\{K, \ldots, K + a - 1\}$, so each of these steps has strictly positive probability. From there it can jump to $K$ as above. Hence *every state can reach* $K$. Each closed communicating class must therefore contain $K$, so there can be at most one such class.

**Step 2: the essential class is positive recurrent.** For $V(i) = i + 1$, Eq. (S2.1) gives a strictly negative drift outside a finite set. The conditional mean elongation is finite because the geometric law has mean $1/p$. The standard Foster drift criterion on a countable state space, together with accessibility of $K$, gives a positive recurrent class containing $K$. The previous step shows this is the only closed class. In particular, its stationary distribution is unique.

**Step 3: the class is aperiodic.** At state $K > a$ the chain can return to $K$ in one step by recruitment with elongation length $a$:

$$P_\theta(K, K) \geq f(K)p(1-p)^{a-1} > 0.$$

Thus the recurrent class contains a self-loop and has period one. Positive recurrence, uniqueness of the essential class, and aperiodicity yield convergence of the law to the unique stationary distribution. □

##### S2.2 Stationary balance and the meaning of a time average

Write $\pi_j = \pi_\theta(j)$ for brevity. The stationary distribution is characterized by

$$\pi_j = \sum_{i\geq 0} \pi_i P_\theta(i,j), \qquad \sum_{j\geq 0} \pi_j = 1. \tag{S2.2}$$

In particular, for a bounded function $h$, the ergodic theorem on the recurrent class gives

$$\frac{1}{T}\sum_{n=0}^{T-1} h(L_n) \xrightarrow[T\to\infty]{\text{a.s.}} \sum_{j\geq 0} h(j)\pi_j. \tag{S2.3}$$

Choosing $h(j) = \mathbf{1}_{\{j=k\}}$ gives the limiting fraction of generations spent at a given length $k$. This statement and convergence of the one-generation distribution are distinct properties, both supported by Proposition S2.1 and related to Eq. (5) of the main article. A sequencing snapshot is modeled as a sample from $\pi_\theta$ only under the additional biological assumption that the population is sufficiently close to stationarity.

#### S3 Structural identifiability from the exact stationary law

This section provides the full proof of Theorem 1 in the main manuscript. Structural identifiability asks whether two parameter sets can generate *exactly the same* stationary telomere length distribution. Theorem 1 addresses this question *conditional on the shortening parameter $a$ being known.* We therefore fix the same integer $a$ when comparing two sets $(L_s, \beta, p)$.

A stationary distribution alone does not reveal a transition matrix. For this reason, we use the probabilities $\pi_0, \pi_1, \ldots$ and the model's stationary-balance equations.

The proof has three explicit steps: (i) determine $p$ from the tail of the stationary distribution, (ii) recover the recruitment probability $f(i)$ at sufficiently large lengths, and (iii) determine $\beta$ and $L_s$ from that recovered recruitment curve.

##### S3.1 Step 1: the exact stationary tail determines $p$

Write $q = 1-p$, where $0 < q < 1$. We first show

$$\boxed{\lim_{j\to\infty} \pi_j^{1/j} = q = 1-p.} \tag{S3.1}$$

The formula says that the rate of exponential decay of the stationary probabilities at very large telomere lengths equals the geometric survival parameter $q$.

**Lemma S3.1** (Geometric decay rate of the stationary tail)**.** *Under the assumptions of Proposition S2.1, Eq.* (S3.1) *holds.*

*Proof.* **Lower bound.** Since $\pi$ is a probability distribution, some state $i_0$ has $\pi_{i_0} > 0$. Recruitment is possible at every length because $f(i_0) > 0$. Therefore, for all $j > r_{i_0}$, the contribution to $\pi_j$ from a recruited elongation starting at $i_0$ is strictly positive:

$$\pi_j \geq \pi_{i_0} f(i_0) p q^{j-r_{i_0}-1}. \tag{S3.2}$$

The prefactor does not depend on $j$. Taking $j$th roots and then letting $j \to \infty$ gives

$$\liminf_{j\to\infty} \pi_j^{1/j} \geq q.$$

**Upper bound.** Fix any real number $t$ satisfying $1 < t < 1/q$. Because the elongation is geometric,

$$\mathbb{E}[t^{Y_n}] = \frac{pt}{1-qt} < \infty. \tag{S3.3}$$

For every $i \geq a$, conditioning on recruitment gives

$$\begin{aligned}\mathbb{E}[t^{L_{n+1}} \mid L_n = i] &= t^{i-a}\left((1-f(i)) + f(i)\frac{pt}{1-qt}\right)\\ &= t^i t^{-a}\left[1 + f(i)\left(\frac{pt}{1-qt} - 1\right)\right].\end{aligned} \tag{S3.4}$$

As $i \to \infty$, the expression in square brackets tends to one, since $f(i) \to 0$. Thus the ratio between the left-hand side and $t^i$ tends to $t^{-a} < 1$. For sufficiently large $i$, there is therefore a number $0 < \rho < 1$ such that

$$\mathbb{E}[t^{L_{n+1}} \mid L_n = i] \leq \rho t^i.$$

For the finitely many remaining states, the conditional moment is finite by Eq. (S3.3). The geometric-drift criterion consequently implies that the positive recurrent stationary distribution has a finite exponential moment:

$$\sum_{j\geq 0} \pi_j t^j < \infty. \tag{S3.5}$$

In particular, $\pi_j t^j \le \sum_{k\ge 0} \pi_k t^k$, so

$$\limsup_{j\to\infty} \pi_j^{1/j} \le 1/t.$$

Since this holds for every $t < 1/q$, sending $t \uparrow 1/q$ yields

$$\limsup_{j\to\infty} \pi_j^{1/j} \le q.$$

The lower bound also yields $\liminf_{j\to\infty} \pi_j^{1/j} \ge q$. Since the upper limit is at most $q$, the ordinary limit exists and equals $q$, proving Eq. (S3.1). □

Therefore, if the infinite stationary probabilities $\pi_j$ were known exactly, then

$$\boxed{p = 1 - \lim_{j\to\infty} \pi_j^{1/j}.} \tag{S3.6}$$

This mathematical identification of $p$ does not use the value of $a$.

#### S3.2 Step 2: recover $f(i)$ from the stationary probabilities

We now fix $a$ and use the known $p$ to reconstruct $f(i)$ for every sufficiently large length. The following identity is the essential calculation.

**Lemma S3.2** (Reconstruction of the recruitment function)**.** *For every integer $i \ge a+1$ such that $\pi_i > 0$, the stationary probabilities satisfy*

$$\boxed{f(i) = 1 - \frac{\pi_{i-a}}{\pi_i} + \frac{p}{\pi_i} \sum_{k=i-a}^{i-1} \pi_k.} \tag{S3.7}$$

*Proof.* For $j \ge 1$, the probability of reaching length $j$ by a recruited elongation, at stationarity, is

$$E_j = p \sum_{i\,:\,r_i \le j-1} \pi_i f(i) q^{j-r_i-1}, \qquad q = 1-p. \tag{S3.8}$$

The probability of reaching $j$ *without* recruitment comes only from the previous length $j+a$, since $j \ge 1$. Consequently, stationary balance reads

$$\pi_j = \pi_{j+a}\big(1 - f(j+a)\big) + E_j. \tag{S3.9}$$

The geometric elongation law has a memoryless tail. Comparing $E_{j+1}$ and $E_j$ in Eq. (S3.8), all contributions already present in $E_j$ acquire one extra factor $q$, and the newly eligible previous length $j+a$ contributes $p\pi_{j+a}f(j+a)$. Hence

$$E_{j+1} = qE_j + p\pi_{j+a}f(j+a). \tag{S3.10}$$

Substitute Eq. (S3.10) into Eq. (S3.9) at $j+1$, and subtract $q$ times Eq. (S3.9) at $j$. This gives, after collecting terms,

$$\begin{aligned}&\pi_{j+a+1}f(j+a+1) - \pi_{j+a}f(j+a)\\ &\quad = \pi_{j+a+1} - \pi_{j+1} - q\big(\pi_{j+a} - \pi_j\big).\end{aligned} \tag{S3.11}$$

Set $i = j+a+1$. The last equation is valid for all $i \geq a+2$ and becomes

$$\begin{aligned}&\pi_i f(i) - \pi_{i-1}f(i-1)\\ &\quad = \pi_i - \pi_{i-a} - q(\pi_{i-1} - \pi_{i-a-1}).\end{aligned} \tag{S3.12}$$

We sum Eq. (S3.12) over indices $i = n+1, n+2, \ldots$, where $n \geq a+1$. Since $\pi_i \to 0$ and $0 < f(i) \leq 1$, the left-hand side telescopes to $-\pi_n f(n)$. The sums on the right-hand side telescope as well:

$$-\pi_n f(n) = -\sum_{k=n-a+1}^{n} \pi_k + q\sum_{k=n-a}^{n-1} \pi_k.$$

Multiplying by $-1$, using $q = 1-p$, and separating the endpoint terms gives

$$\pi_n f(n) = \pi_n - \pi_{n-a} + p\sum_{k=n-a}^{n-1} \pi_k. \tag{S3.13}$$

Dividing by $\pi_n > 0$ proves Eq. (S3.7). □

##### S3.3 Step 3: recover $\beta$ and $L_s$

For every $i > L_s$, the specific recruitment formula implies

$$\frac{1}{f(i)} - 1 = \beta(i - L_s). \tag{S3.14}$$

The *left-hand side* is now determined from $\pi$ through Eq. (S3.7), once $a$ and $p$ are known. The *right-hand side* is an affine function of the integer $i$, with slope $\beta$ and intercept $-\beta L_s$.

Choose any two distinct sufficiently large lengths $i_1 < i_2$ such that both are above $L_s$ and their stationary probabilities are positive. Let

$$h_1 = \frac{1}{f(i_1)} - 1, \qquad h_2 = \frac{1}{f(i_2)} - 1.$$

Equation (S3.14) gives $h_1 = \beta(i_1 - L_s)$ and $h_2 = \beta(i_2 - L_s)$. Subtracting the two equations and then solving yields

$$\boxed{\beta = \frac{h_2 - h_1}{i_2 - i_1}, \qquad L_s = i_1 - \frac{h_1}{\beta}.} \tag{S3.15}$$

#### S3.4 Complete proof of the identifiability theorem in the main article

**Theorem S3.3** (Proof of main Theorem 1: identifiability conditional on shortening)**.** *Consider the untruncated telomere length model in Eqs.* (S1.1)–(S1.4)*, with fixed and known integer* $a \geq 1$*, finite integer* $L_s \geq 0$*,* $\beta > 0$*, and* $0 < p < 1$*. If two admissible parameter triples* $(L_s, \beta, p)$ *generate exactly the same stationary distribution, then the triples are equal. In other words, for known* $a$*, the stationary distribution uniquely determines* $(L_s, \beta, p)$*.*

*Proof.* Suppose that $(a, L_s^{(1)}, \beta_1, p_1)$ and $(a, L_s^{(2)}, \beta_2, p_2)$ generate the same exact stationary probabilities $\pi_j$.

First, $p_1 = p_2$. By Lemma S3.1, the common probabilities satisfy

$$1 - p_1 = \lim_{j\to\infty} \pi_j^{1/j} = 1 - p_2.$$

Thus both parameter vectors have the same geometric elongation parameter $p$.

Second, their recruitment functions coincide at all sufficiently large lengths. The integer $a$ and the elongation parameter $p$ are the same. Equation (S3.7) therefore computes the same value of $f(i)$ from the common stationary probabilities for every sufficiently large $i$.

Third, $\beta_1 = \beta_2$ and $L_s^{(1)} = L_s^{(2)}$. Select two different integers $i_1, i_2$ larger than both threshold values and large enough for the reconstruction formula to be valid. The recovered values $f(i_1)$ and $f(i_2)$ agree for both parameter sets. By Eq. (S3.15), the slope $\beta$ and threshold $L_s$ are then uniquely determined. Thus $\beta_1 = \beta_2$ and $L_s^{(1)} = L_s^{(2)}$.

All three coordinates agree, proving the theorem. $\square$

##### S3.5 Implications and limitations of the identifiability theorem for $a$

The theorem fixes $a$ *before* comparing parameter vectors. It does not establish injectivity of the full four-parameter mapping $\theta \mapsto \pi_\theta$ when distinct values of $a$ are allowed. In particular, the fact that the stationary law identifies $p$ does not prove that it also identifies $a$; the reconstruction formula changes with $a$.

For a proposed integer $a$, Eq. (S3.7) constructs a candidate recruitment function from the exact stationary distribution. To establish full identifiability without knowing $a$, one would additionally need to prove that only one integer $a$ makes that candidate a valid function of the prescribed form, with the correct behavior at the short-length boundary. Such a uniqueness argument across different values of $a$ is *not supplied here.* We therefore make no claim that an exact equality across different $a$ values exists, nor that it is impossible.

Numerically, scanning candidate values of $a$ and plotting the optimized $W_1$ profile is a sensible selection procedure. A lowest observed value of $W_1$ for one $a$ is evidence about fit quality for that dataset and search procedure.

### S4 Finite-state stationary distribution used for fitting

##### S4.1 Truncation and conservation of probability mass

Write $M = L_{\max}$ and use the finite state space $\{0, 1, \ldots, M\}$. One probability-conserving convention is to combine all jumps that would reach or exceed $M$ at the boundary state $M$. With $r_i = (i-a)_+$, this defines

$$P_\theta^{(M)}(i,j) = \begin{cases} (1 - f(i))\mathbf{1}_{\{j=r_i\}} + f(i)pq^{j-r_i-1}\mathbf{1}_{\{j \geq r_i+1\}}, & j < M, \\ f(i)q^{M-r_i-1}, & j = M, \end{cases} \tag{S4.1}$$

where $q = 1-p$ and $0 \leq i \leq M$. The final entry is the exact geometric probability of $Y_n \geq M - r_i$, including jumps exceeding $M$. Since $r_i \leq M-1$, this expression is well defined and each row sums to one. Explicitly,

$$\sum_{j=0}^{M} P_\theta^{(M)}(i,j) = (1 - f(i)) + f(i)\left[\sum_{k=1}^{M-r_i-1} pq^{k-1} + q^{M-r_i-1}\right] = 1. \tag{S4.2}$$

This is an exact description of the kernel.

#### S4.2 Ergodicity of the truncated Markov chain

**Proposition S4.1** (Ergodicity of the truncated chain)**.** *For the truncated transition kernel in Eq.* (S4.1)*, with $M \geq 1$, $a \geq 1$, $0 < p < 1$, and $f(i) > 0$, there exists a unique stationary distribution $\pi_\theta^{(M)}$. Moreover, for any initial probability distribution $\nu$,*

$$\lim_{n\to\infty} \nu(P_\theta^{(M)})^n = \pi_\theta^{(M)}.$$

*Proof.* Let $q = 1-p$. Every state $i \in \{0, \ldots, M\}$ can reach $M$ in one step with positive probability:

$$P_\theta^{(M)}(i, M) = f(i) q^{M-r_i-1} > 0.$$

Hence, the chain has a unique closed communicating class containing $M$. Since $P_\theta^{(M)}(M, M) > 0$, this class is aperiodic. The result follows from the ergodic theorem for finite-state Markov chains. □

#### S4.3 Spectral interpretation and a qualification

For the matrix, the unique stationary eigenvalue is 1 and all other eigenvalues have modulus smaller than 1. Let $|\lambda_2|$ denote the second-largest eigenvalue modulus. This quantity describes the asymptotic spectral convergence rate. For a matrix that is diagonalizable, a bound of the form $C|\lambda_2|^n$ can be obtained; for a non-diagonalizable matrix, Jordan blocks can introduce a polynomial factor in $n$. Without assuming diagonalizability, a valid general statement is that for every $r$ satisfying $|\lambda_2| < r < 1$, there is a constant $C_r$ such that

$$\|\nu(P_\theta^{(M)})^n - \pi_\theta^{(M)}\|_{\mathrm{TV}} \leq C_r r^n. \tag{S4.3}$$

Thus a positive spectral gap is a convergence diagnostic, but its use as an *exact* $C|\lambda_2|^n$ bound needs an additional matrix assumption.

### S5 Wasserstein distance and finite sampling

#### S5.1 Exact discrete one-dimensional expression

For probability distributions $\pi$ and $\mu$ on $\{0, \ldots, M\}$, define their cumulative probabilities by $F_\pi(j) = \sum_{k=0}^{j} \pi_k$ and $F_\mu(j) = \sum_{k=0}^{j} \mu_k$. The one-dimensional 1-Wasserstein distance, with cost equal to the absolute length difference in bp, is

$$\boxed{W_1(\pi, \mu) = \sum_{j=0}^{M-1} |F_\pi(j) - F_\mu(j)|.} \tag{S5.1}$$

#### S5.2 Proof of the $N^{-1/2}$ fluctuation bound

The following argument proves Proposition 1 (and the bound in Eq. (8)) of the main manuscript.

**Proposition S5.1** (Expected empirical Wasserstein fluctuation)**.** *Let $X_1, \ldots, X_N$ be independent samples from a probability distribution $\pi$ on $\{0, \ldots, M\}$. Let $\widehat{\pi}_N$ be their empirical distribution and $F(j) = \sum_{k=0}^{j} \pi_k$. Then*

$$\boxed{\mathbb{E}[W_1(\widehat{\pi}_N, \pi)] \leq \frac{1}{\sqrt{N}} \sum_{j=0}^{M-1} \sqrt{F(j)(1 - F(j))}.} \tag{S5.2}$$

*Proof.* For each fixed integer $j$, the empirical cumulative distribution is the average of $N$ indicator variables:

$$\widehat{F}_N(j) = \frac{1}{N} \sum_{m=1}^{N} \mathbf{1}_{\{X_m \leq j\}}. \tag{S5.3}$$

The indicators are independent Bernoulli variables with success probability $F(j)$. Consequently,

$$\mathbb{E}[\widehat{F}_N(j)] = F(j), \qquad \mathrm{Var}(\widehat{F}_N(j)) = \frac{F(j)(1 - F(j))}{N}. \tag{S5.4}$$

By the Cauchy–Schwarz inequality, for every random variable with finite variance,

$$\mathbb{E}\left[\left|\widehat{F}_N(j) - F(j)\right|\right] \leq \sqrt{\mathbb{E}[(\widehat{F}_N(j) - F(j))^2]} \tag{S5.5}$$

$$= \sqrt{\frac{F(j)(1 - F(j))}{N}}. \tag{S5.6}$$

Finally insert this bound into the exact sum for $W_1$ in Eq. (S5.1) and use linearity of expectation:

$$\begin{aligned}\mathbb{E}[W_1(\widehat{\pi}_N, \pi)] &= \sum_{j=0}^{M-1} \mathbb{E}[|\widehat{F}_N(j) - F(j)|] \\ &\leq \frac{1}{\sqrt{N}} \sum_{j=0}^{M-1} \sqrt{F(j)(1 - F(j))}.\end{aligned}$$

This proves the proposition. □

The bound controls the *expected* Wasserstein fluctuation, not every realization. The constant depends on the underlying distribution and need not be the same for a short-telomere and a long-telomere strain. Supplementary Figure S1 provides a reproducible Monte Carlo illustration based on an explicitly defined synthetic model, rather than an unprovided experimental dataset.

##### S5.3 Structural versus practical identifiability

Even when two different parameter sets produce distinct stationary laws, those laws may be difficult to distinguish using only $N$ observations. The bound above makes the relevant sampling scale explicit. If the distance between two predicted stationary distributions is much smaller than the characteristic sampling fluctuations for the available $N$, several distinct parameter combinations may fit the same empirical histogram. This is *practical* difficulty, not a contradiction of the theorem in Section S3. Conversely, finding several optimizer runs with similar Wasserstein distances is not sufficient to demonstrate an exact equality of stationary laws.

#### S6 Bootstrap comparison and interpretation

##### S6.1 Definition used in the main manuscript

For an experimental analysis unit with $N$ filtered observed telomere lengths, the manuscript specifies a nonparametric bootstrap with $B = 1000$ resamples. Each bootstrap dataset contains $N$ observations sampled with replacement from the observed lengths. Let $\pi_{\text{exp}}$ be the original empirical distribution, $\widehat{\pi}_N^{(b)}$ the empirical distribution of bootstrap replicate $b$, and

$$T_b = W_1(\widehat{\pi}_N^{(b)}, \pi_{\text{exp}}), \qquad b = 1, \dots, B. \tag{S6.1}$$

The bootstrap mean and sample standard deviation are

$$\overline{T} = \frac{1}{B}\sum_{b=1}^{B} T_b, \tag{S6.2}$$

$$s_T = \sqrt{\frac{1}{B-1}\sum_{b=1}^{B}(T_b - \overline{T})^2}. \tag{S6.3}$$

The threshold is

$$\boxed{W_{\text{thr}} = \overline{T} + 3s_T.} \tag{S6.4}$$

For a fitted parameter vector $\widehat{\theta}$ and its predicted stationary distribution $\pi_{\widehat{\theta}}$, the comparison is

$$W_1(\pi_{\text{exp}}, \pi_{\widehat{\theta}}) \le W_{\text{thr}}. \tag{S6.5}$$

Fits satisfying this rule are called *compatible with the estimated sampling scale* in the main analysis. A fit above the reference is not automatically a useless estimate: it indicates a remaining discrepancy larger than that particular empirical sampling reference.

### S7 Numerical parameter estimation

#### S7.1 Objective and fixed-$a$ parameter scan

The study estimates the parameters by minimizing the Wasserstein distance between $\pi_{\text{exp}}$ and the numerically computed stationary distribution:

$$\widehat{\theta} \in \arg\min_{\theta\in\Theta} W_1(\pi_{\text{exp}}, \pi_\theta). \tag{S7.1}$$

Since $a$ is integer valued, it is treated by a profile search rather than included as a continuous coordinate in CMA-ES. For each $a \in \{2, 3, \ldots, 20\}$, the remaining parameters $(\beta, L_s, p)$ are optimized, with $L_s$ restricted to integer values when constructing the Markov chain. The profile value at fixed $a$ is the lowest Wasserstein distance obtained among the corresponding optimization runs. The best fit over all tested $a$ values is the smallest profile value. The profile is useful both for choosing an estimate and for displaying alternatives with similar fit quality. It is not a mathematical proof that the full parameter vector is globally identifiable.

#### S7.2 Search ranges and preprocessing dependence

The numerical domain reported in the supplied manuscript is

$$10^{-4} \leq \beta \leq 0.20, \qquad 0.003 \leq p \leq 0.20, \qquad 0 \leq L_s \leq L_s^{\max}, \tag{S7.2}$$

where the selected upper bound for experimental fits is

$$L_s^{\max} = \min\{400, \lfloor Q_{0.01}(L_{\exp}) \rfloor\}. \tag{S7.3}$$

Here $Q_{0.01}$ denotes the first percentile of the empirical lengths after preprocessing. This is an *inference restriction* motivated by the observed short-length range. As in the main article, the set of admissible $L_s$ values is integer valued even if a continuous optimizer temporarily proposes non-integer coordinates.

#### S7.3 CMA-ES runs and computational budget

The manuscript specifies 15 independent CMA-ES initializations at every fixed $a$, a population size of 12 and at most 100 generations. The total number of independent CMA-ES runs per empirical distribution is therefore $19 \times 15 = 285$. A single run evaluates at most approximately $12 \times 100 = 1200$ candidate parameter vectors, and the nominal upper budget per distribution is

$$19 \times 15 \times 12 \times 100 = 342\,000 \tag{S7.4}$$

objective evaluations, without counting possible early stopping or additional evaluations performed by a specific implementation. The conventional population-size heuristic for $d = 3$ parameters is $4 + \lfloor 3 \log d \rfloor = 7$; the selected size 12 is a deliberate enlargement of this heuristic, not a mathematical requirement. Multiple random starts reduce the risk that a poor local solution is mistaken for the best fit, but they do not guarantee global optimality.

#### S7.4 Synthetic distributions

For a specified parameter vector $\theta^{\star}$, synthetic validation begins by calculating a stationary distribution from the Markov model, drawing independent length observations from that distribution, and applying the same parameter-estimation procedure as for experimental samples. The known generating distribution and parameters provide two separate targets: agreement of the fitted *distribution* with the true law, and agreement of the fitted *parameters* with $\theta^\star$. Good distributional agreement need not imply precise recovery of each coordinate when $N$ is small.

### Part II. Supplementary Figures

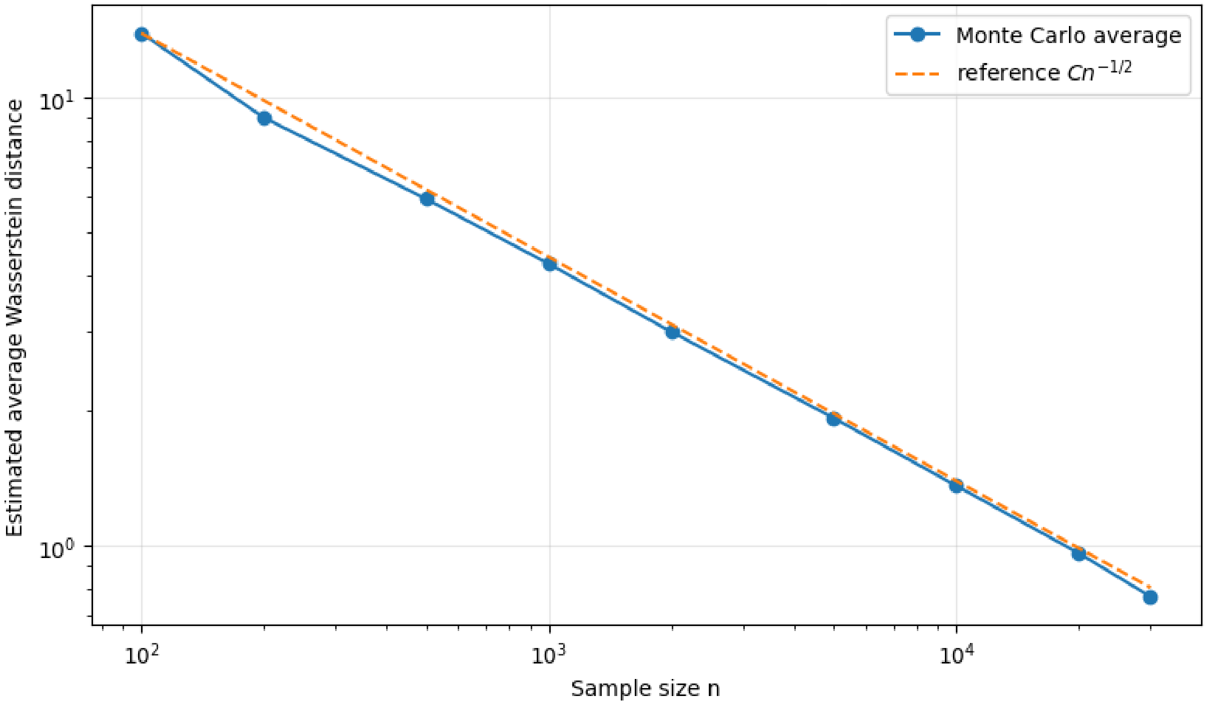


**Supplementary Figure S1. Log–log plot of the Monte Carlo estimate of $\mathbb{E}[W_1(\widehat{\pi}_n, \pi)]$ as a function of the sample size $n$.** The dashed reference line has slope $-1/2$, corresponding to the $n^{-1/2}$ rate of Proposition S5.1. The numerical curve is nearly parallel to this reference, confirming the theoretical prediction.

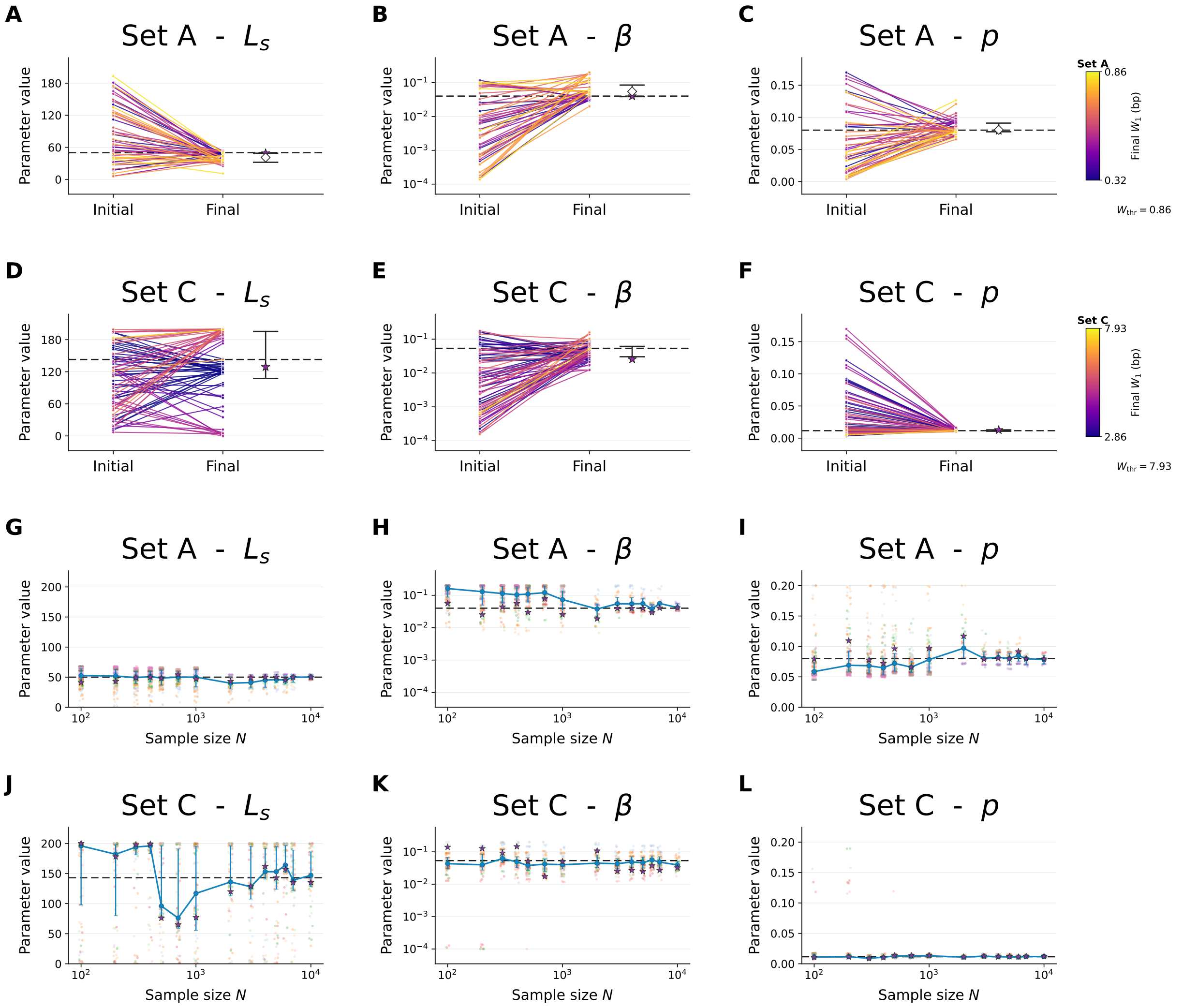


**Supplementary Figure S2. Parameter inference performance using synthetic data.** Synthetic data were generated for Set A with ground-truth parameters $(a, \beta, L_s, p) = (8, 0.040, 50, 0.080)$ and for Set C with ground-truth parameters $(a, \beta, L_s, p) = (8, 0.054, 143, 0.012)$. **(A–C)** Initial and final estimates of $L_s$, $\beta$, and $p$, respectively, for Set A, as in Fig. 3A–C. **(D–F)** Same as panels A–C for Set C. **(G–I)** Parameter estimates of $L_s$, $\beta$, and $p$, respectively, as a function of sample size $N$ for Set A, as in Fig. 3E–G. **(J–L)** Same as panels G–I for Set C.

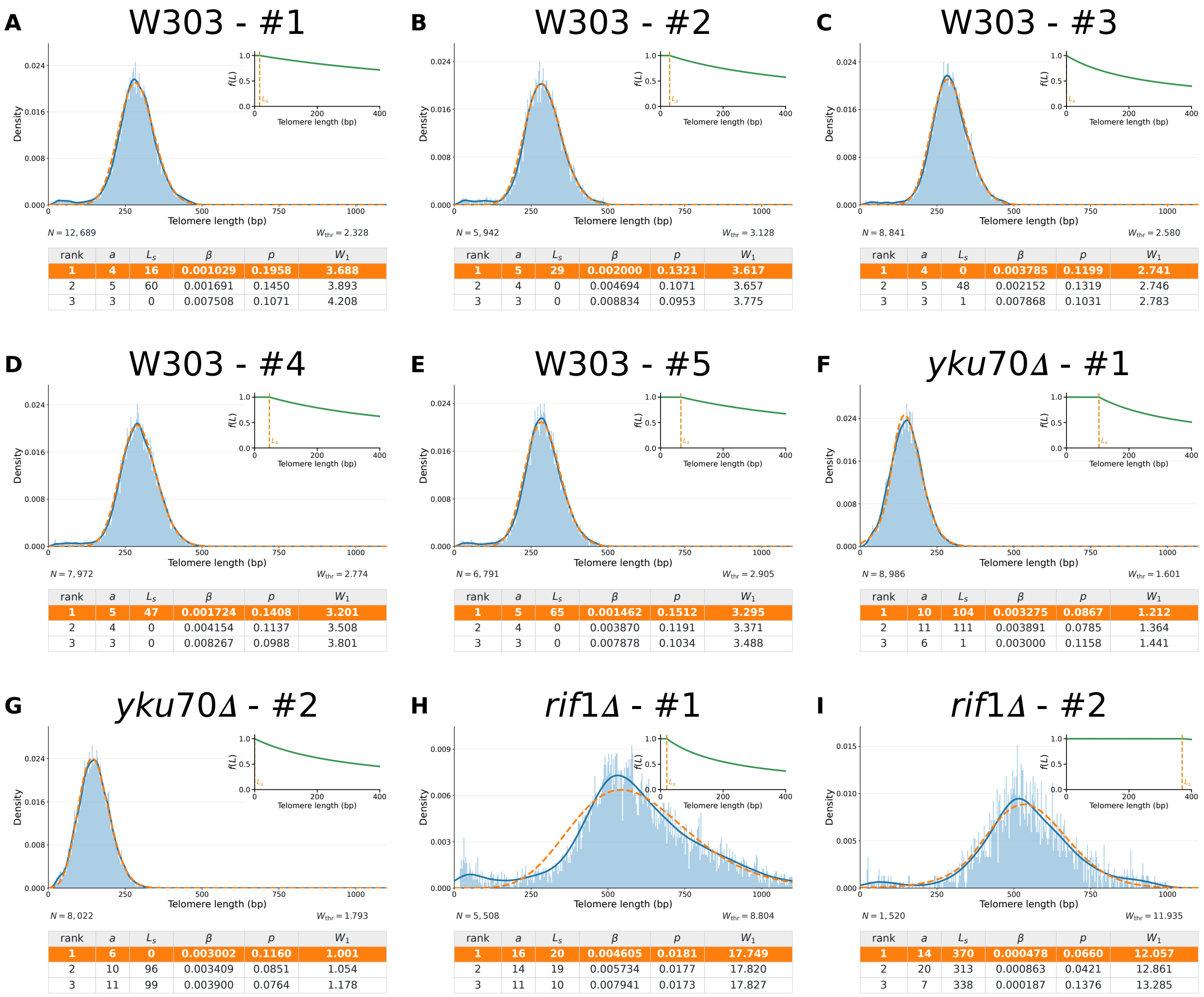


**A** W303 - #1 ($N = 12,689$, $W_{thr} = 2.328$)

| rank | $a$ | $L_s$ | $\beta$ | $p$ | $W_1$ |
|---|---|---|---|---|---|
| **1** | **4** | **16** | **0.001029** | **0.1958** | **3.688** |
| 2 | 5 | 60 | 0.001691 | 0.1450 | 3.893 |
| 3 | 3 | 0 | 0.007508 | 0.1071 | 4.208 |

**B** W303 - #2 ($N = 5,942$, $W_{thr} = 3.128$)

| rank | $a$ | $L_s$ | $\beta$ | $p$ | $W_1$ |
|---|---|---|---|---|---|
| **1** | **5** | **29** | **0.002000** | **0.1321** | **3.617** |
| 2 | 4 | 0 | 0.004694 | 0.1071 | 3.657 |
| 3 | 3 | 0 | 0.008834 | 0.0953 | 3.775 |

**C** W303 - #3 ($N = 8,841$, $W_{thr} = 2.580$)

| rank | $a$ | $L_s$ | $\beta$ | $p$ | $W_1$ |
|---|---|---|---|---|---|
| **1** | **4** | **0** | **0.003785** | **0.1199** | **2.741** |
| 2 | 5 | 48 | 0.002152 | 0.1319 | 2.746 |
| 3 | 3 | 1 | 0.007868 | 0.1031 | 2.783 |

**D** W303 - #4 ($N = 7,972$, $W_{thr} = 2.774$)

| rank | $a$ | $L_s$ | $\beta$ | $p$ | $W_1$ |
|---|---|---|---|---|---|
| **1** | **5** | **47** | **0.001724** | **0.1408** | **3.201** |
| 2 | 4 | 0 | 0.004154 | 0.1137 | 3.508 |
| 3 | 3 | 0 | 0.008267 | 0.0988 | 3.801 |

**E** W303 - #5 ($N = 6,791$, $W_{thr} = 2.905$)

| rank | $a$ | $L_s$ | $\beta$ | $p$ | $W_1$ |
|---|---|---|---|---|---|
| **1** | **5** | **65** | **0.001462** | **0.1512** | **3.295** |
| 2 | 4 | 0 | 0.003870 | 0.1191 | 3.371 |
| 3 | 3 | 0 | 0.007878 | 0.1034 | 3.488 |

**F** *yku70Δ* - #1 ($N = 8,986$, $W_{thr} = 1.601$)

| rank | $a$ | $L_s$ | $\beta$ | $p$ | $W_1$ |
|---|---|---|---|---|---|
| **1** | **10** | **104** | **0.003275** | **0.0867** | **1.212** |
| 2 | 11 | 111 | 0.003891 | 0.0785 | 1.364 |
| 3 | 6 | 1 | 0.003000 | 0.1158 | 1.441 |

**G** *yku70Δ* - #2 ($N = 8,022$, $W_{thr} = 1.793$)

| rank | $a$ | $L_s$ | $\beta$ | $p$ | $W_1$ |
|---|---|---|---|---|---|
| **1** | **6** | **0** | **0.003002** | **0.1160** | **1.001** |
| 2 | 10 | 96 | 0.003409 | 0.0851 | 1.054 |
| 3 | 11 | 99 | 0.003900 | 0.0764 | 1.178 |

**H** *rif1Δ* - #1 ($N = 5,508$, $W_{thr} = 8.804$)

| rank | $a$ | $L_s$ | $\beta$ | $p$ | $W_1$ |
|---|---|---|---|---|---|
| **1** | **16** | **20** | **0.004605** | **0.0181** | **17.749** |
| 2 | 14 | 19 | 0.005734 | 0.0177 | 17.820 |
| 3 | 11 | 10 | 0.007941 | 0.0173 | 17.827 |

**I** *rif1Δ* - #2 ($N = 1,520$, $W_{thr} = 11.935$)

| rank | $a$ | $L_s$ | $\beta$ | $p$ | $W_1$ |
|---|---|---|---|---|---|
| **1** | **14** | **370** | **0.000478** | **0.0660** | **12.057** |
| 2 | 20 | 313 | 0.000863 | 0.0421 | 12.861 |
| 3 | 7 | 338 | 0.000187 | 0.1376 | 13.285 |

**Supplementary Figure S3. Estimation of parameters from experimental telomere length distributions in independent replicates. (A–I)** Fits of telomere length distributions from wild-type laboratory strain W303 **(A–E)** and its mutant derivatives *yku70*Δ **(F and G)** and *rif1*Δ **(H and I)** for the indicated independent replicates.

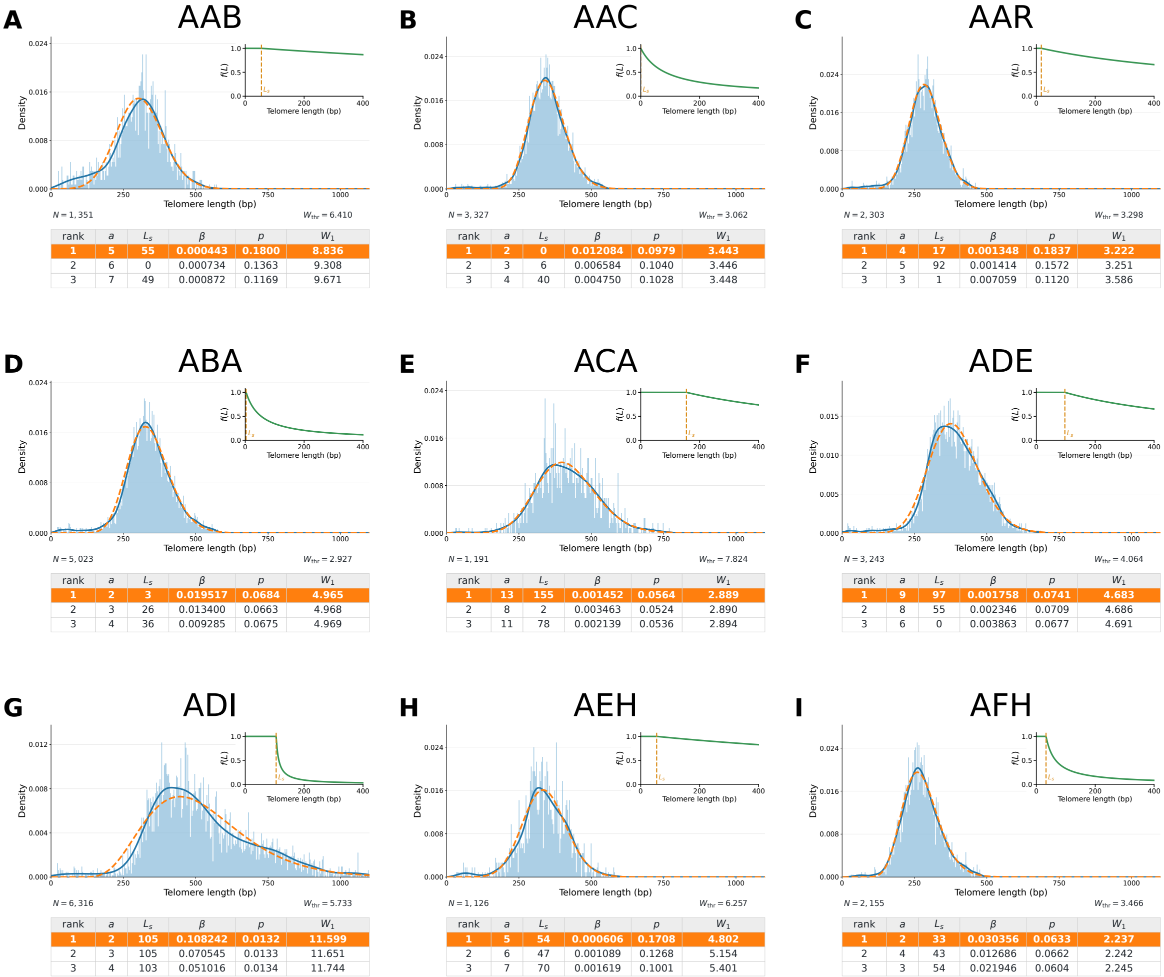


**A AAB**

| rank | $a$ | $L_s$ | $\beta$ | $p$ | $W_1$ |
|---|---|---|---|---|---|
| 1 | 5 | 55 | 0.000443 | 0.1800 | 8.836 |
| 2 | 6 | 0 | 0.000734 | 0.1363 | 9.308 |
| 3 | 7 | 49 | 0.000872 | 0.1169 | 9.671 |

**B AAC**

| rank | $a$ | $L_s$ | $\beta$ | $p$ | $W_1$ |
|---|---|---|---|---|---|
| 1 | 2 | 0 | 0.012084 | 0.0979 | 3.443 |
| 2 | 3 | 6 | 0.006584 | 0.1040 | 3.446 |
| 3 | 4 | 40 | 0.004750 | 0.1028 | 3.448 |

**C AAR**

| rank | $a$ | $L_s$ | $\beta$ | $p$ | $W_1$ |
|---|---|---|---|---|---|
| 1 | 4 | 17 | 0.001348 | 0.1837 | 3.222 |
| 2 | 5 | 92 | 0.001414 | 0.1572 | 3.251 |
| 3 | 3 | 1 | 0.007059 | 0.1120 | 3.586 |

**D ABA**

| rank | $a$ | $L_s$ | $\beta$ | $p$ | $W_1$ |
|---|---|---|---|---|---|
| 1 | 2 | 3 | 0.019517 | 0.0684 | 4.965 |
| 2 | 3 | 26 | 0.013400 | 0.0663 | 4.968 |
| 3 | 4 | 36 | 0.009285 | 0.0675 | 4.969 |

**E ACA**

| rank | $a$ | $L_s$ | $\beta$ | $p$ | $W_1$ |
|---|---|---|---|---|---|
| 1 | 13 | 155 | 0.001452 | 0.0564 | 2.889 |
| 2 | 8 | 2 | 0.003463 | 0.0524 | 2.890 |
| 3 | 11 | 78 | 0.002139 | 0.0536 | 2.894 |

**F ADE**

| rank | $a$ | $L_s$ | $\beta$ | $p$ | $W_1$ |
|---|---|---|---|---|---|
| 1 | 9 | 97 | 0.001758 | 0.0741 | 4.683 |
| 2 | 8 | 55 | 0.002346 | 0.0709 | 4.686 |
| 3 | 6 | 0 | 0.003863 | 0.0677 | 4.691 |

**G ADI**

| rank | $a$ | $L_s$ | $\beta$ | $p$ | $W_1$ |
|---|---|---|---|---|---|
| 1 | 2 | 105 | 0.108242 | 0.0132 | 11.599 |
| 2 | 3 | 105 | 0.070545 | 0.0133 | 11.651 |
| 3 | 4 | 103 | 0.051016 | 0.0134 | 11.744 |

**H AEH**

| rank | $a$ | $L_s$ | $\beta$ | $p$ | $W_1$ |
|---|---|---|---|---|---|
| 1 | 5 | 54 | 0.000606 | 0.1708 | 4.802 |
| 2 | 6 | 47 | 0.001089 | 0.1268 | 5.154 |
| 3 | 7 | 70 | 0.001619 | 0.1001 | 5.401 |

**I AFH**

| rank | $a$ | $L_s$ | $\beta$ | $p$ | $W_1$ |
|---|---|---|---|---|---|
| 1 | 2 | 33 | 0.030356 | 0.0633 | 2.237 |
| 2 | 4 | 43 | 0.012686 | 0.0662 | 2.242 |
| 3 | 3 | 54 | 0.021946 | 0.0604 | 2.245 |

**Supplementary Figure S4. Estimation of parameters from telomere length distributions of natural yeast isolates (Part 1).** (A–I) Same as Fig. 5D–F with the indicated strains.

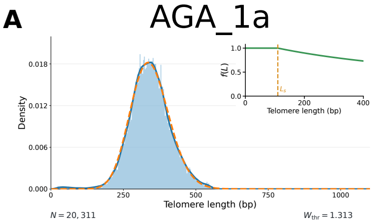


| rank | $a$ | $L_s$ | $\beta$ | $p$ | $W_1$ |
|---|---|---|---|---|---|
| **1** | **6** | **110** | **0.001268** | **0.1292** | **2.278** |
| 2 | 7 | 163 | 0.001332 | 0.1158 | 2.280 |
| 3 | 5 | 20 | 0.001577 | 0.1332 | 2.282 |

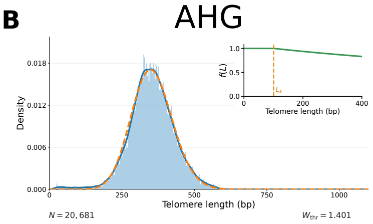


| rank | $a$ | $L_s$ | $\beta$ | $p$ | $W_1$ |
|---|---|---|---|---|---|
| **1** | **5** | **101** | **0.000678** | **0.1710** | **2.980** |
| 2 | 6 | 120 | 0.000964 | 0.1363 | 3.015 |
| 3 | 7 | 117 | 0.001678 | 0.1028 | 3.241 |

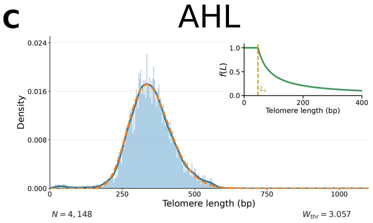


| rank | $a$ | $L_s$ | $\beta$ | $p$ | $W_1$ |
|---|---|---|---|---|---|
| **1** | **2** | **47** | **0.024603** | **0.0618** | **2.888** |
| 2 | 4 | 68 | 0.011218 | 0.0623 | 2.893 |
| 3 | 5 | 82 | 0.008677 | 0.0623 | 2.894 |

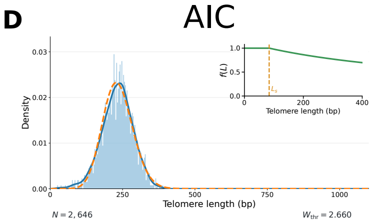


| rank | $a$ | $L_s$ | $\beta$ | $p$ | $W_1$ |
|---|---|---|---|---|---|
| **1** | **5** | **84** | **0.001371** | **0.1665** | **3.410** |
| 2 | 4 | 7 | 0.001712 | 0.1807 | 3.412 |
| 3 | 6 | 84 | 0.002208 | 0.1262 | 3.965 |

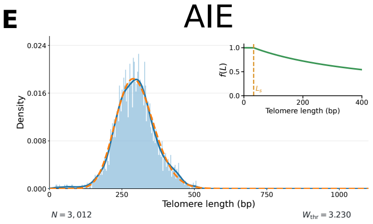


| rank | $a$ | $L_s$ | $\beta$ | $p$ | $W_1$ |
|---|---|---|---|---|---|
| **1** | **6** | **33** | **0.002287** | **0.1047** | **2.378** |
| 2 | 7 | 84 | 0.002008 | 0.1008 | 2.388 |
| 3 | 5 | 2 | 0.003946 | 0.0936 | 2.395 |

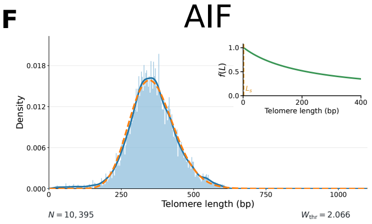


| rank | $a$ | $L_s$ | $\beta$ | $p$ | $W_1$ |
|---|---|---|---|---|---|
| **1** | **5** | **2** | **0.004655** | **0.0765** | **3.527** |
| 2 | 7 | 55 | 0.002094 | 0.0882 | 3.531 |
| 3 | 6 | 2 | 0.002271 | 0.0930 | 3.531 |

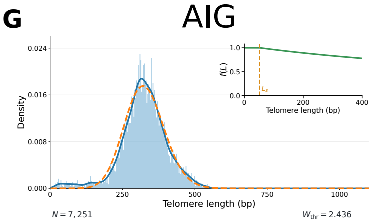


| rank | $a$ | $L_s$ | $\beta$ | $p$ | $W_1$ |
|---|---|---|---|---|---|
| **1** | **5** | **52** | **0.000814** | **0.1638** | **6.382** |
| 2 | 6 | 44 | 0.001630 | 0.1147 | 6.513 |
| 3 | 4 | 3 | 0.006404 | 0.0828 | 6.774 |

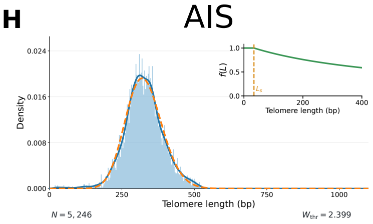


| rank | $a$ | $L_s$ | $\beta$ | $p$ | $W_1$ |
|---|---|---|---|---|---|
| **1** | **5** | **34** | **0.001907** | **0.1291** | **3.122** |
| 2 | 6 | 141 | 0.001206 | 0.1367 | 3.128 |
| 3 | 4 | 16 | 0.004797 | 0.1017 | 3.132 |

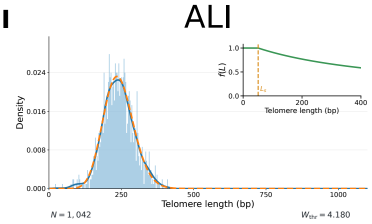


| rank | $a$ | $L_s$ | $\beta$ | $p$ | $W_1$ |
|---|---|---|---|---|---|
| **1** | **5** | **51** | **0.002023** | **0.1452** | **1.544** |
| 2 | 6 | 99 | 0.002039 | 0.1300 | 1.547 |
| 3 | 4 | 2 | 0.004618 | 0.1202 | 1.551 |

**Supplementary Figure S5. Estimation of parameters from telomere length distributions of natural yeast isolates (Part 2).** (A–I) Same as Fig. 5D–F with the indicated strains.

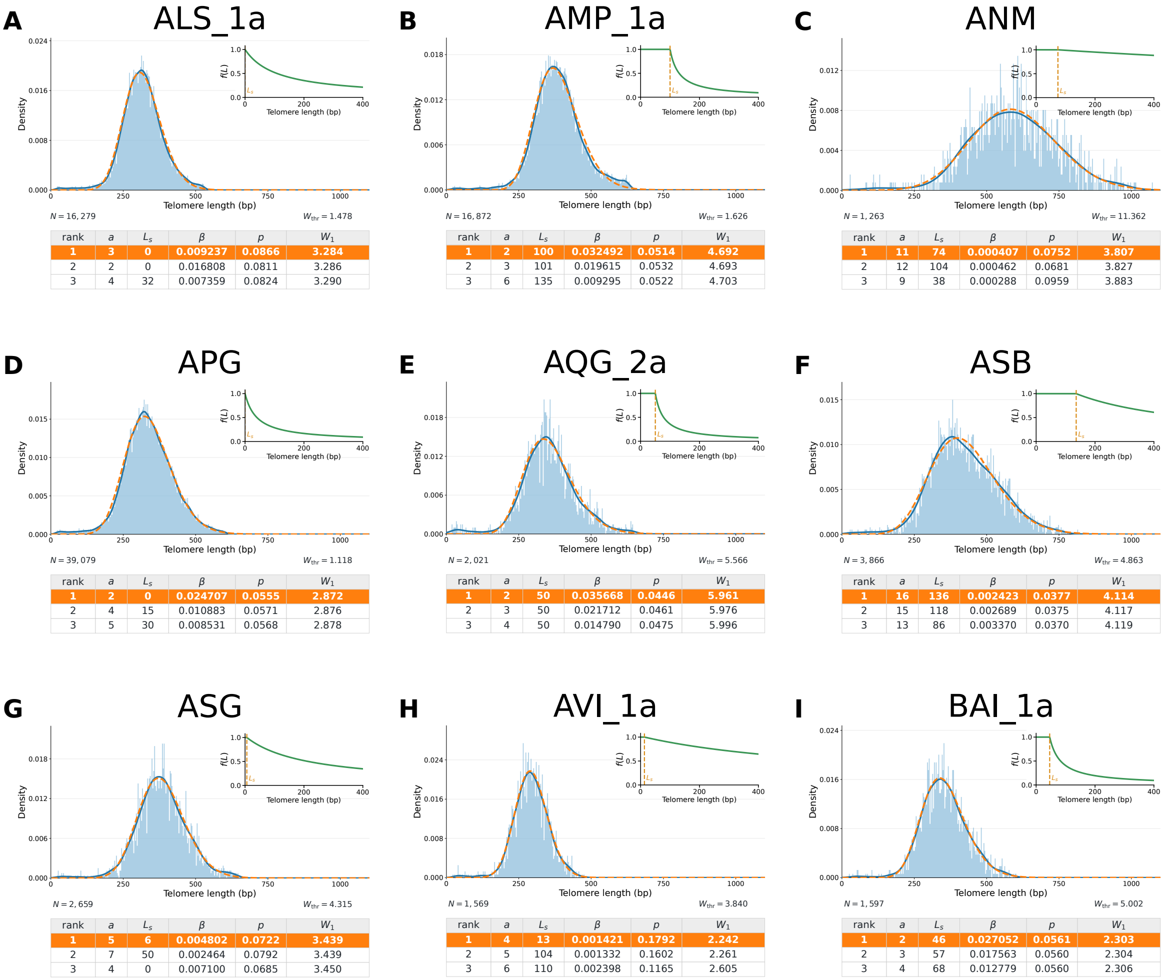


ALS_1a

| rank | $a$ | $L_s$ | $\beta$ | $p$ | $W_1$ |
|---|---|---|---|---|---|
| 1 | 3 | 0 | 0.009237 | 0.0866 | 3.284 |
| 2 | 2 | 0 | 0.016808 | 0.0811 | 3.286 |
| 3 | 4 | 32 | 0.007359 | 0.0824 | 3.290 |

AMP_1a

| rank | $a$ | $L_s$ | $\beta$ | $p$ | $W_1$ |
|---|---|---|---|---|---|
| 1 | 2 | 100 | 0.032492 | 0.0514 | 4.692 |
| 2 | 3 | 101 | 0.019615 | 0.0532 | 4.693 |
| 3 | 6 | 135 | 0.009295 | 0.0522 | 4.703 |

ANM

| rank | $a$ | $L_s$ | $\beta$ | $p$ | $W_1$ |
|---|---|---|---|---|---|
| 1 | 11 | 74 | 0.000407 | 0.0752 | 3.807 |
| 2 | 12 | 104 | 0.000462 | 0.0681 | 3.827 |
| 3 | 9 | 38 | 0.000288 | 0.0959 | 3.883 |

APG

| rank | $a$ | $L_s$ | $\beta$ | $p$ | $W_1$ |
|---|---|---|---|---|---|
| 1 | 2 | 0 | 0.024707 | 0.0555 | 2.872 |
| 2 | 4 | 15 | 0.010883 | 0.0571 | 2.876 |
| 3 | 5 | 30 | 0.008531 | 0.0568 | 2.878 |

AQG_2a

| rank | $a$ | $L_s$ | $\beta$ | $p$ | $W_1$ |
|---|---|---|---|---|---|
| 1 | 2 | 50 | 0.035668 | 0.0446 | 5.961 |
| 2 | 3 | 50 | 0.021712 | 0.0461 | 5.976 |
| 3 | 4 | 50 | 0.014790 | 0.0475 | 5.996 |

ASB

| rank | $a$ | $L_s$ | $\beta$ | $p$ | $W_1$ |
|---|---|---|---|---|---|
| 1 | 16 | 136 | 0.002423 | 0.0377 | 4.114 |
| 2 | 15 | 118 | 0.002689 | 0.0375 | 4.117 |
| 3 | 13 | 86 | 0.003370 | 0.0370 | 4.119 |

ASG

| rank | $a$ | $L_s$ | $\beta$ | $p$ | $W_1$ |
|---|---|---|---|---|---|
| 1 | 5 | 6 | 0.004802 | 0.0722 | 3.439 |
| 2 | 7 | 50 | 0.002464 | 0.0792 | 3.439 |
| 3 | 4 | 0 | 0.007100 | 0.0685 | 3.450 |

AVI_1a

| rank | $a$ | $L_s$ | $\beta$ | $p$ | $W_1$ |
|---|---|---|---|---|---|
| 1 | 4 | 13 | 0.001421 | 0.1792 | 2.242 |
| 2 | 5 | 104 | 0.001332 | 0.1602 | 2.261 |
| 3 | 6 | 110 | 0.002398 | 0.1165 | 2.605 |

BAI_1a

| rank | $a$ | $L_s$ | $\beta$ | $p$ | $W_1$ |
|---|---|---|---|---|---|
| 1 | 2 | 46 | 0.027052 | 0.0561 | 2.303 |
| 2 | 3 | 57 | 0.017563 | 0.0560 | 2.304 |
| 3 | 4 | 68 | 0.012779 | 0.0560 | 2.306 |

**Supplementary Figure S6. Estimation of parameters from telomere length distributions of natural yeast isolates (Part 3).** (A–I) Same as Fig. 5D–F with the indicated strains.

**A** BAK_1a

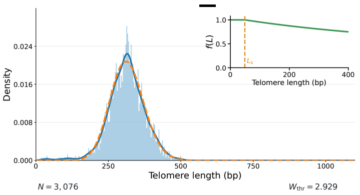


$N = 3,076$ $W_{thr} = 2.929$

| rank | $a$ | $L_s$ | $\beta$ | $p$ | $W_1$ |
|---|---|---|---|---|---|
| **1** | **4** | **49** | **0.000951** | **0.1999** | **3.199** |
| 2 | 5 | 93 | 0.001491 | 0.1509 | 3.308 |
| 3 | 3 | 8 | 0.006707 | 0.1107 | 3.540 |

**B** BAL_1a

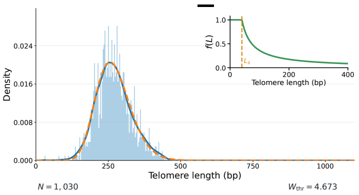


$N = 1,030$ $W_{thr} = 4.673$

| rank | $a$ | $L_s$ | $\beta$ | $p$ | $W_1$ |
|---|---|---|---|---|---|
| **1** | **2** | **40** | **0.029612** | **0.0674** | **1.758** |
| 2 | 3 | 61 | 0.021632 | 0.0638 | 1.758 |
| 3 | 7 | 98 | 0.007134 | 0.0665 | 1.764 |

**C** BBF

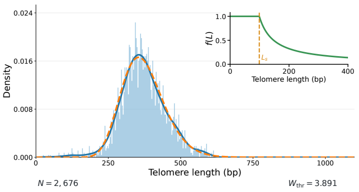


$N = 2,676$ $W_{thr} = 3.891$

| rank | $a$ | $L_s$ | $\beta$ | $p$ | $W_1$ |
|---|---|---|---|---|---|
| **1** | **3** | **99** | **0.020484** | **0.0533** | **3.085** |
| 2 | 4 | 105 | 0.014432 | 0.0540 | 3.086 |
| 3 | 5 | 113 | 0.010907 | 0.0546 | 3.089 |

**D** BBM_1a

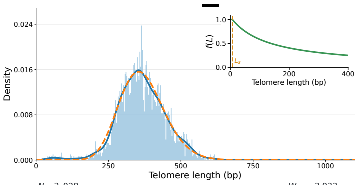


$N = 3,028$ $W_{thr} = 3.933$

| rank | $a$ | $L_s$ | $\beta$ | $p$ | $W_1$ |
|---|---|---|---|---|---|
| **1** | **4** | **7** | **0.007647** | **0.0682** | **3.438** |
| 2 | 7 | 67 | 0.003527 | 0.0705 | 3.443 |
| 3 | 5 | 11 | 0.005102 | 0.0723 | 3.444 |

**E** BCE_3a

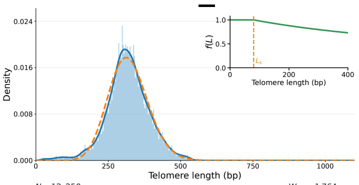


$N = 12,250$ $W_{thr} = 1.764$

| rank | $a$ | $L_s$ | $\beta$ | $p$ | $W_1$ |
|---|---|---|---|---|---|
| **1** | **6** | **80** | **0.001154** | **0.1311** | **4.205** |
| 2 | 5 | 15 | 0.000982 | 0.1544 | 4.211 |
| 3 | 7 | 95 | 0.001747 | 0.1034 | 4.236 |

**F** BDC_1c

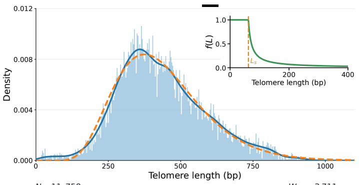


$N = 11,758$ $W_{thr} = 3.711$

| rank | $a$ | $L_s$ | $\beta$ | $p$ | $W_1$ |
|---|---|---|---|---|---|
| **1** | **2** | **62** | **0.096376** | **0.0160** | **5.146** |
| 2 | 3 | 66 | 0.064141 | 0.0160 | 5.147 |
| 3 | 4 | 68 | 0.047249 | 0.0161 | 5.147 |

**G** BDF_1d

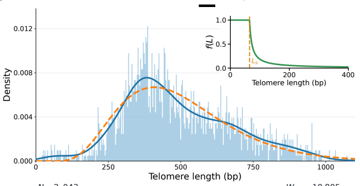


$N = 2,043$ $W_{thr} = 10.805$

| rank | $a$ | $L_s$ | $\beta$ | $p$ | $W_1$ |
|---|---|---|---|---|---|
| **1** | **2** | **65** | **0.126209** | **0.0113** | **13.056** |
| 2 | 3 | 67 | 0.083341 | 0.0113 | 13.057 |
| 3 | 4 | 70 | 0.062336 | 0.0113 | 13.058 |

**H** BDH_1b

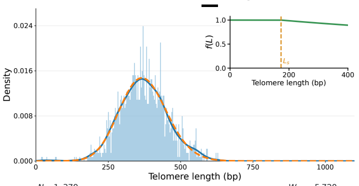


$N = 1,379$ $W_{thr} = 5.720$

| rank | $a$ | $L_s$ | $\beta$ | $p$ | $W_1$ |
|---|---|---|---|---|---|
| **1** | **6** | **173** | **0.000524** | **0.1511** | **1.941** |
| 2 | 5 | 188 | 0.000384 | 0.1869 | 1.946 |
| 3 | 7 | 25 | 0.001819 | 0.0881 | 2.247 |

**I** BDM_1a

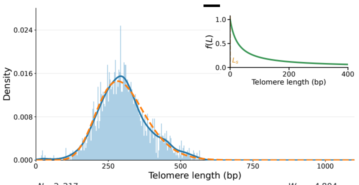


$N = 2,217$ $W_{thr} = 4.804$

| rank | $a$ | $L_s$ | $\beta$ | $p$ | $W_1$ |
|---|---|---|---|---|---|
| **1** | **2** | **0** | **0.037068** | **0.0435** | **4.151** |
| 2 | 5 | 23 | 0.013703 | 0.0437 | 4.156 |
| 3 | 4 | 2 | 0.015769 | 0.0458 | 4.158 |

277

**Supplementary Figure S7. Estimation of parameters from telomere length distributions of natural yeast isolates (Part 4).** (A–I) Same as Fig. 5D–F with the indicated strains.

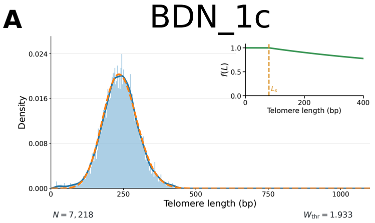


| rank | $a$ | $L_s$ | $\beta$ | $p$ | $W_1$ |
|---|---|---|---|---|---|
| **1** | **5** | **80** | **0.000894** | **0.1753** | **1.727** |
| 2 | 6 | 66 | 0.001465 | 0.1334 | 2.046 |
| 3 | 7 | 81 | 0.002069 | 0.1083 | 2.450 |

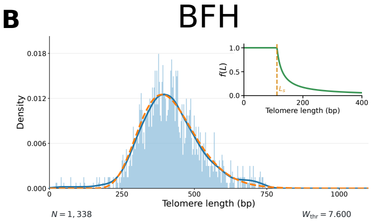


| rank | $a$ | $L_s$ | $\beta$ | $p$ | $W_1$ |
|---|---|---|---|---|---|
| **1** | **2** | **112** | **0.054041** | **0.0313** | **5.059** |
| 2 | 4 | 122 | 0.025995 | 0.0314 | 5.062 |
| 3 | 5 | 125 | 0.019949 | 0.0318 | 5.063 |

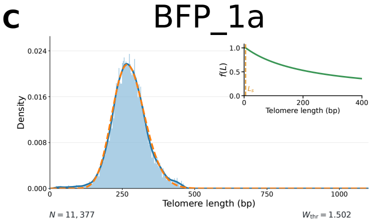


| rank | $a$ | $L_s$ | $\beta$ | $p$ | $W_1$ |
|---|---|---|---|---|---|
| **1** | **4** | **5** | **0.004566** | **0.1134** | **2.325** |
| 2 | 5 | 41 | 0.002689 | 0.1238 | 2.326 |
| 3 | 6 | 94 | 0.002170 | 0.1206 | 2.336 |

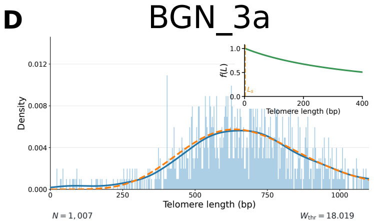


| rank | $a$ | $L_s$ | $\beta$ | $p$ | $W_1$ |
|---|---|---|---|---|---|
| **1** | **20** | **2** | **0.002440** | **0.0194** | **9.329** |
| 2 | 19 | 2 | 0.002780 | 0.0189 | 9.415 |
| 3 | 18 | 6 | 0.003188 | 0.0183 | 9.535 |

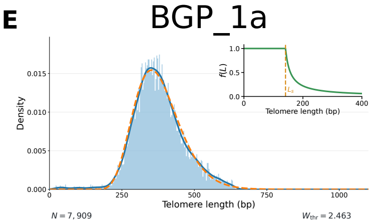


| rank | $a$ | $L_s$ | $\beta$ | $p$ | $W_1$ |
|---|---|---|---|---|---|
| **1** | **2** | **141** | **0.059856** | **0.0366** | **3.718** |
| 2 | 3 | 146 | 0.039227 | 0.0366 | 3.720 |
| 3 | 4 | 150 | 0.028470 | 0.0368 | 3.721 |

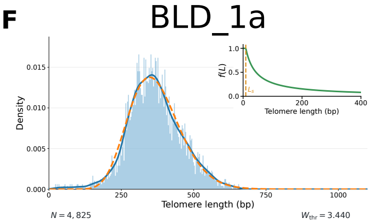


| rank | $a$ | $L_s$ | $\beta$ | $p$ | $W_1$ |
|---|---|---|---|---|---|
| **1** | **2** | **9** | **0.028697** | **0.0466** | **3.624** |
| 2 | 5 | 31 | 0.009957 | 0.0479 | 3.630 |
| 3 | 6 | 56 | 0.009005 | 0.0457 | 3.631 |

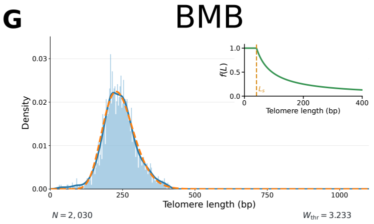


| rank | $a$ | $L_s$ | $\beta$ | $p$ | $W_1$ |
|---|---|---|---|---|---|
| **1** | **3** | **41** | **0.018863** | **0.0733** | **2.742** |
| 2 | 2 | 43 | 0.033926 | 0.0687 | 2.745 |
| 3 | 4 | 43 | 0.011776 | 0.0780 | 2.745 |

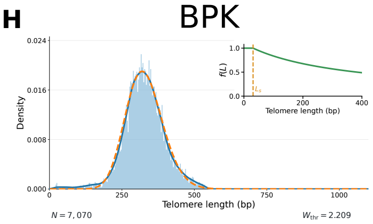


| rank | $a$ | $L_s$ | $\beta$ | $p$ | $W_1$ |
|---|---|---|---|---|---|
| **1** | **5** | **31** | **0.002840** | **0.1095** | **3.776** |
| 2 | 4 | 5 | 0.004701 | 0.1006 | 3.783 |
| 3 | 6 | 86 | 0.001913 | 0.1147 | 3.786 |

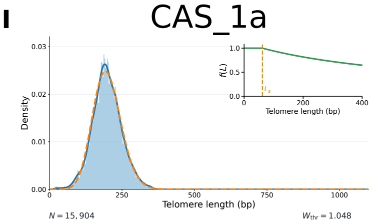


| rank | $a$ | $L_s$ | $\beta$ | $p$ | $W_1$ |
|---|---|---|---|---|---|
| **1** | **5** | **62** | **0.001623** | **0.1644** | **1.328** |
| 2 | 6 | 75 | 0.002395 | 0.1296 | 1.465 |
| 3 | 4 | 7 | 0.001557 | 0.1932 | 1.502 |

**Supplementary Figure S8. Estimation of parameters from telomere length distributions of natural yeast isolates (Part 5).** (A–I) Same as Fig. 5D–F with the indicated strains.

**A** CCN

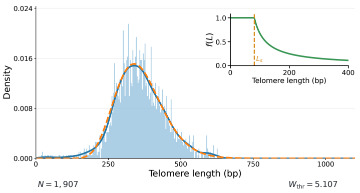


$N = 1,907$ $W_{thr} = 5.107$

| rank | $a$ | $L_s$ | $\beta$ | $p$ | $W_1$ |
|---|---|---|---|---|---|
| **1** | **3** | **81** | **0.025443** | **0.0439** | **3.086** |
| 2 | 4 | 89 | 0.018680 | 0.0439 | 3.087 |
| 3 | 6 | 102 | 0.011433 | 0.0446 | 3.091 |

**B** CDG_1a

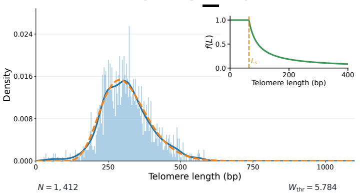


$N = 1,412$ $W_{thr} = 5.784$

| rank | $a$ | $L_s$ | $\beta$ | $p$ | $W_1$ |
|---|---|---|---|---|---|
| **1** | **3** | **64** | **0.032842** | **0.0395** | **3.543** |
| 2 | 2 | 61 | 0.051548 | 0.0391 | 3.544 |
| 3 | 4 | 66 | 0.023167 | 0.0402 | 3.544 |

**C** CDN_1a

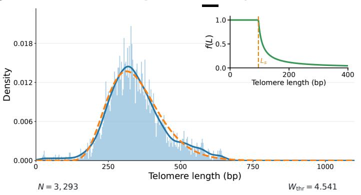


$N = 3,293$ $W_{thr} = 4.541$

| rank | $a$ | $L_s$ | $\beta$ | $p$ | $W_1$ |
|---|---|---|---|---|---|
| **1** | **2** | **96** | **0.072572** | **0.0297** | **6.528** |
| 2 | 3 | 96 | 0.045700 | 0.0303 | 6.565 |
| 3 | 4 | 96 | 0.032539 | 0.0307 | 6.605 |

**D** CEL_1a

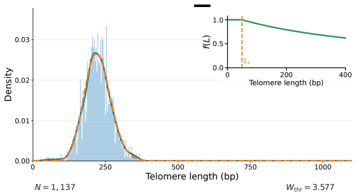


$N = 1,137$ $W_{thr} = 3.577$

| rank | $a$ | $L_s$ | $\beta$ | $p$ | $W_1$ |
|---|---|---|---|---|---|
| **1** | **4** | **49** | **0.001743** | **0.1924** | **1.515** |
| 2 | 5 | 97 | 0.002077 | 0.1592 | 1.643 |
| 3 | 3 | 7 | 0.007011 | 0.1342 | 1.723 |

**E** CEQ_1a

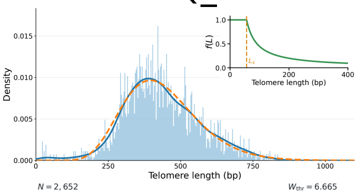


$N = 2,652$ $W_{thr} = 6.665$

| rank | $a$ | $L_s$ | $\beta$ | $p$ | $W_1$ |
|---|---|---|---|---|---|
| **1** | **4** | **56** | **0.028206** | **0.0235** | **5.950** |
| 2 | 5 | 59 | 0.021988 | 0.0237 | 5.952 |
| 3 | 7 | 71 | 0.015421 | 0.0236 | 5.952 |

**F** CFS

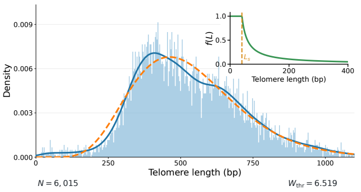


$N = 6,015$ $W_{thr} = 6.519$

| rank | $a$ | $L_s$ | $\beta$ | $p$ | $W_1$ |
|---|---|---|---|---|---|
| **1** | **3** | **40** | **0.053209** | **0.0142** | **7.624** |
| 2 | 4 | 43 | 0.039425 | 0.0142 | 7.624 |
| 3 | 2 | 42 | 0.082342 | 0.0141 | 7.624 |

**G** CGH_1

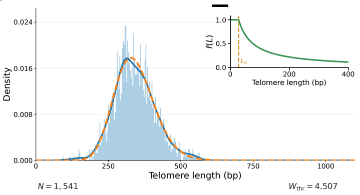


$N = 1,541$ $W_{thr} = 4.507$

| rank | $a$ | $L_s$ | $\beta$ | $p$ | $W_1$ |
|---|---|---|---|---|---|
| **1** | **2** | **28** | **0.020471** | **0.0699** | **1.754** |
| 2 | 5 | 68 | 0.006831 | 0.0716 | 1.759 |
| 3 | 6 | 89 | 0.005630 | 0.0706 | 1.765 |

**H** CGH_3

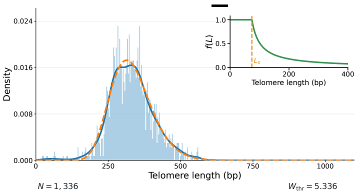


$N = 1,336$ $W_{thr} = 5.336$

| rank | $a$ | $L_s$ | $\beta$ | $p$ | $W_1$ |
|---|---|---|---|---|---|
| **1** | **2** | **74** | **0.035871** | **0.0521** | **3.139** |
| 2 | 4 | 86 | 0.016204 | 0.0531 | 3.143 |
| 3 | 3 | 88 | 0.024608 | 0.0508 | 3.144 |

**I** CKB

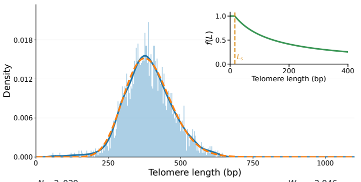


$N = 3,039$ $W_{thr} = 3.946$

| rank | $a$ | $L_s$ | $\beta$ | $p$ | $W_1$ |
|---|---|---|---|---|---|
| **1** | **4** | **16** | **0.007645** | **0.0665** | **2.544** |
| 2 | 5 | 30 | 0.005600 | 0.0678 | 2.547 |
| 3 | 6 | 53 | 0.004500 | 0.0676 | 2.549 |

**Supplementary Figure S9. Estimation of parameters from telomere length distributions of natural yeast isolates (Part 6).** (A–I) Same as Fig. 5D–F with the indicated strains.

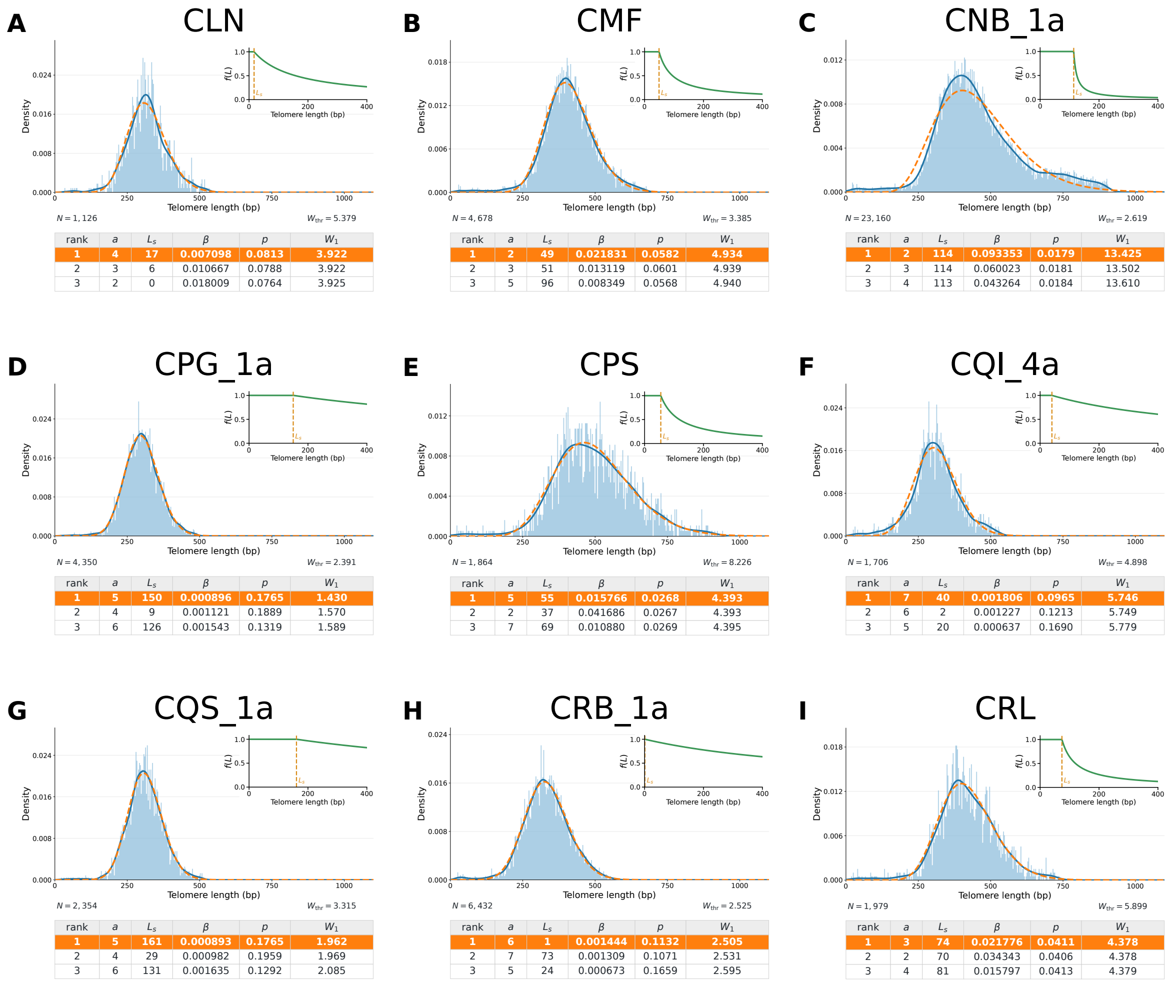


**Supplementary Figure S10. Estimation of parameters from telomere length distributions of natural yeast isolates (Part 7).** (A–I) Same as Fig. 5D–F with the indicated strains.

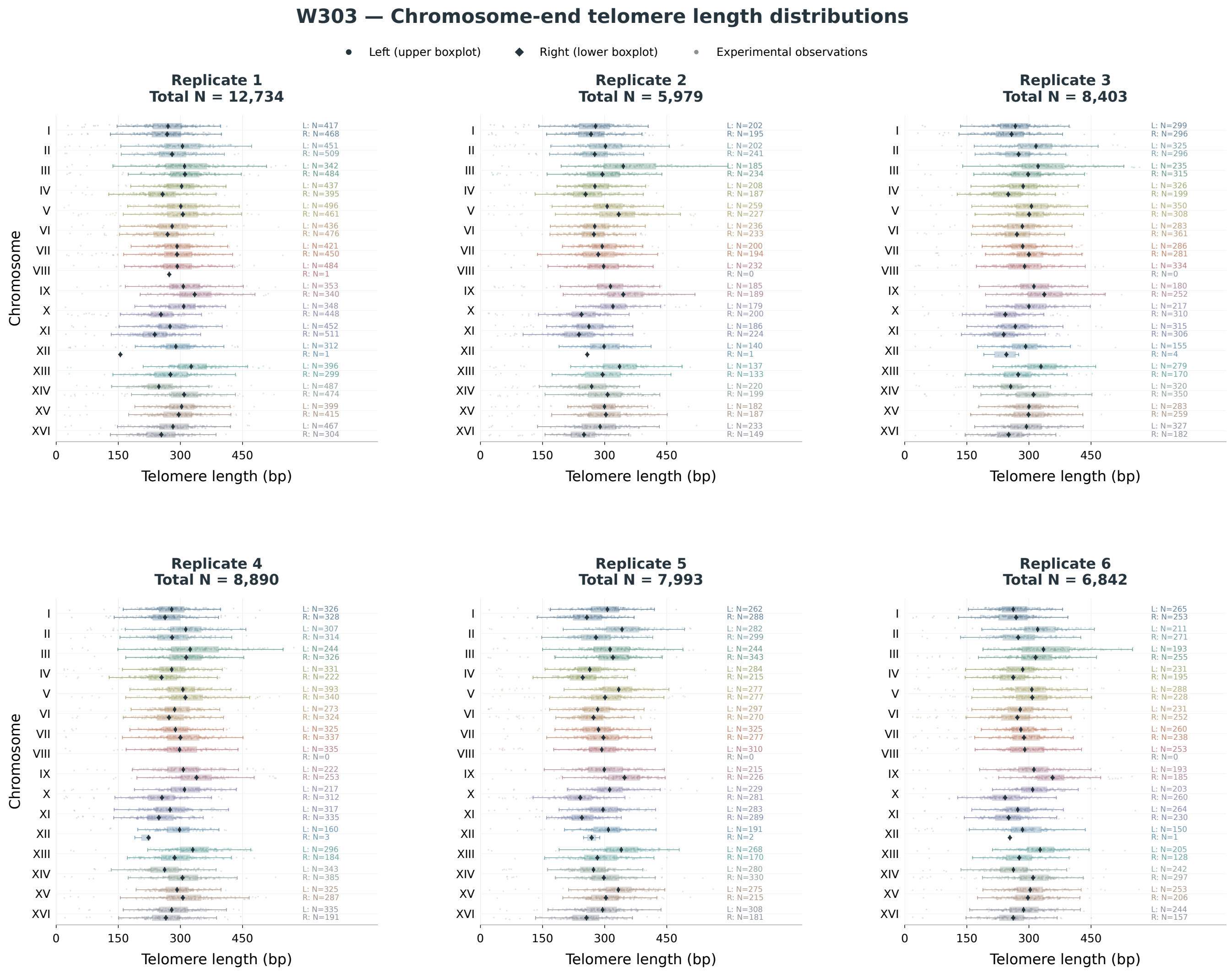


**Supplementary Figure S11. Chromosome-end-specific telomere length distributions across biological replicates of wild-type W303 strain.** Experimental telomere length distributions across the 32 chromosomes ends for the indicated replicate, represented as boxplots. The sample sizes (N) are indicated.

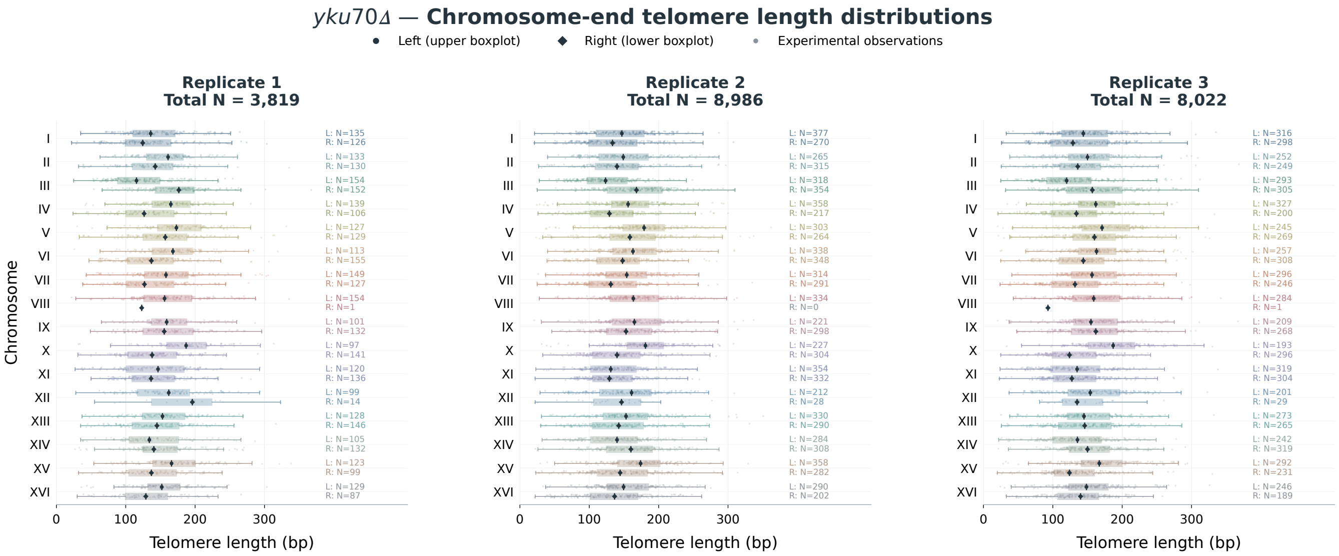


**Supplementary Figure S12. Chromosome-end-specific telomere length distributions across biological replicates of the *yku70Δ* mutant.** Experimental telomere length distributions across the 32 chromosomes ends for the indicated replicate, represented as boxplots. The sample sizes (N) are indicated.

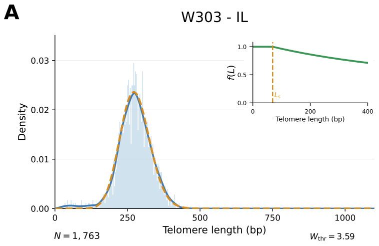


| rank | $a$ | $L_s$ | $\beta$ | $p$ | $W_1$ |
|---|---|---|---|---|---|
| **1** | **4** | **69** | **0.001220** | **0.2000** | **3.263** |
| 2 | 2 | 0 | 0.012739 | 0.1128 | 3.743 |
| 3 | 3 | 22 | 0.007894 | 0.1127 | 3.755 |

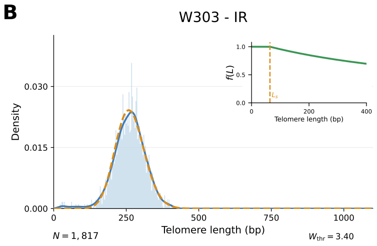


| rank | $a$ | $L_s$ | $\beta$ | $p$ | $W_1$ |
|---|---|---|---|---|---|
| **1** | **4** | **64** | **0.001295** | **0.1993** | **3.456** |
| 2 | 3 | 1 | 0.005963 | 0.1316 | 4.070 |
| 3 | 5 | 71 | 0.002635 | 0.1338 | 4.262 |

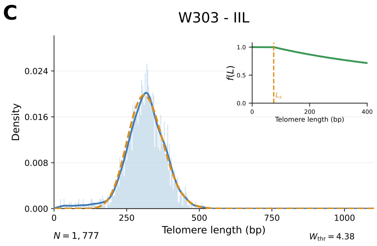


| rank | $a$ | $L_s$ | $\beta$ | $p$ | $W_1$ |
|---|---|---|---|---|---|
| **1** | **5** | **75** | **0.001221** | **0.1550** | **5.185** |
| 2 | 4 | 0 | 0.003702 | 0.1162 | 5.609 |
| 3 | 3 | 0 | 0.007537 | 0.1001 | 5.899 |

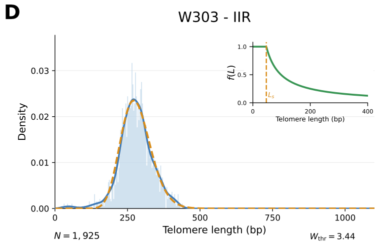


| rank | $a$ | $L_s$ | $\beta$ | $p$ | $W_1$ |
|---|---|---|---|---|---|
| **1** | **2** | **47** | **0.019570** | **0.0926** | **3.553** |
| 2 | 4 | 80 | 0.008946 | 0.0918 | 3.558 |
| 3 | 5 | 90 | 0.005545 | 0.0990 | 3.562 |

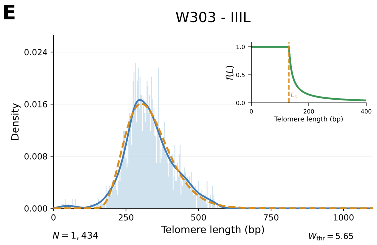


| rank | $a$ | $L_s$ | $\beta$ | $p$ | $W_1$ |
|---|---|---|---|---|---|
| **1** | **2** | **131** | **0.082996** | **0.0326** | **4.795** |
| 2 | 3 | 131 | 0.052114 | 0.0332 | 4.797 |
| 3 | 4 | 138 | 0.039892 | 0.0327 | 4.797 |

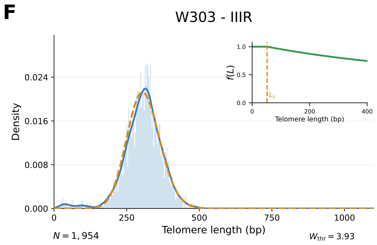


| rank | $a$ | $L_s$ | $\beta$ | $p$ | $W_1$ |
|---|---|---|---|---|---|
| **1** | **4** | **52** | **0.000988** | **0.1998** | **5.127** |
| 2 | 3 | 3 | 0.005943 | 0.1196 | 5.904 |
| 3 | 2 | 0 | 0.011896 | 0.1088 | 6.093 |

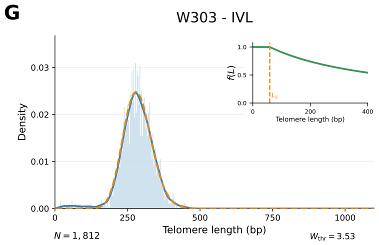


| rank | $a$ | $L_s$ | $\beta$ | $p$ | $W_1$ |
|---|---|---|---|---|---|
| **1** | **4** | **59** | **0.002483** | **0.1611** | **3.392** |
| 2 | 3 | 2 | 0.003897 | 0.1598 | 3.393 |
| 3 | 2 | 11 | 0.010866 | 0.1279 | 3.642 |

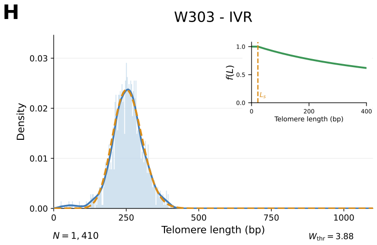


| rank | $a$ | $L_s$ | $\beta$ | $p$ | $W_1$ |
|---|---|---|---|---|---|
| **1** | **4** | **22** | **0.001632** | **0.1816** | **3.048** |
| 2 | 5 | 71 | 0.002449 | 0.1386 | 3.207 |
| 3 | 3 | 12 | 0.007944 | 0.1154 | 3.363 |

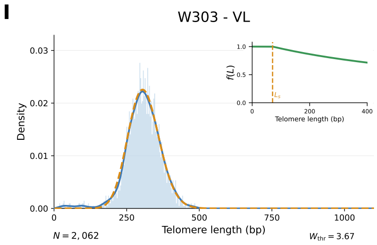


| rank | $a$ | $L_s$ | $\beta$ | $p$ | $W_1$ |
|---|---|---|---|---|---|
| **1** | **4** | **71** | **0.001212** | **0.1944** | **3.309** |
| 2 | 3 | 0 | 0.004678 | 0.1375 | 3.673 |
| 3 | 2 | 5 | 0.010843 | 0.1182 | 3.962 |

**Supplementary Figure S13. Wild-type W303 chromosome-end-specific telomere length distributions fitted by our model (Part 1). (A–I)** Each graph shows the best fit, as in Fig. 5A, for the indicated chromosome end. The table summarizes the three best fits. The probability of telomerase recruitment, $f(L)$, is shown for the best-fit parameters $L_s$ and $\beta$.

A

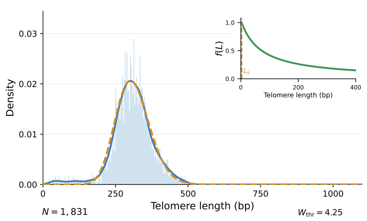


| rank | $a$ | $L_s$ | $\beta$ | $p$ | $W_1$ |
|---|---|---|---|---|---|
| **1** | **2** | **3** | **0.014259** | **0.0950** | **5.270** |
| 2 | 3 | 18 | 0.008518 | 0.0973 | 5.272 |
| 3 | 4 | 44 | 0.006179 | 0.0961 | 5.277 |

B

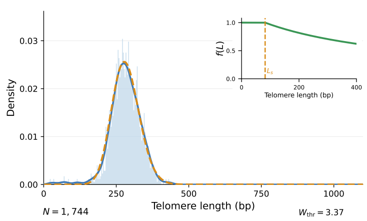


| rank | $a$ | $L_s$ | $\beta$ | $p$ | $W_1$ |
|---|---|---|---|---|---|
| **1** | **4** | **82** | **0.001903** | **0.1817** | **2.945** |
| 2 | 3 | 25 | 0.004729 | 0.1516 | 3.050 |
| 3 | 2 | 12 | 0.010032 | 0.1366 | 3.162 |

C

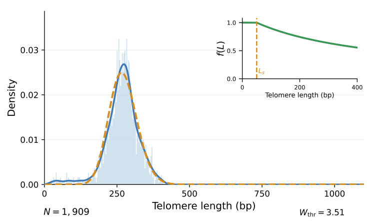


| rank | $a$ | $L_s$ | $\beta$ | $p$ | $W_1$ |
|---|---|---|---|---|---|
| **1** | **4** | **50** | **0.002286** | **0.1669** | **5.870** |
| 2 | 3 | 3 | 0.004753 | 0.1479 | 5.956 |
| 3 | 2 | 0 | 0.010811 | 0.1293 | 6.096 |

D

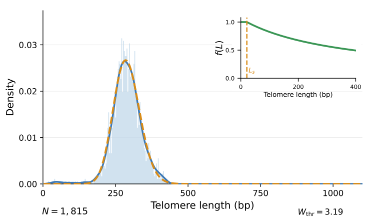


| rank | $a$ | $L_s$ | $\beta$ | $p$ | $W_1$ |
|---|---|---|---|---|---|
| **1** | **3** | **21** | **0.002705** | **0.1941** | **2.819** |
| 2 | 4 | 111 | 0.001872 | 0.1881 | 2.832 |
| 3 | 2 | 21 | 0.009015 | 0.1481 | 2.847 |

E

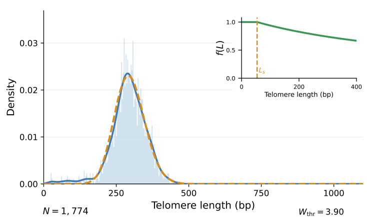


| rank | $a$ | $L_s$ | $\beta$ | $p$ | $W_1$ |
|---|---|---|---|---|---|
| **1** | **4** | **54** | **0.001441** | **0.1858** | **4.855** |
| 2 | 3 | 0 | 0.004829 | 0.1384 | 5.141 |
| 3 | 2 | 0 | 0.010723 | 0.1215 | 5.311 |

F

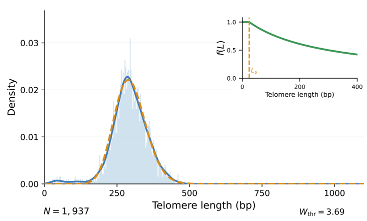


| rank | $a$ | $L_s$ | $\beta$ | $p$ | $W_1$ |
|---|---|---|---|---|---|
| **1** | **4** | **24** | **0.003640** | **0.1269** | **4.016** |
| 2 | 3 | 2 | 0.006456 | 0.1168 | 4.044 |
| 3 | 2 | 0 | 0.012761 | 0.1069 | 4.113 |

G

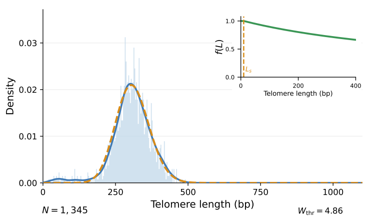


| rank | $a$ | $L_s$ | $\beta$ | $p$ | $W_1$ |
|---|---|---|---|---|---|
| **1** | **4** | **10** | **0.001294** | **0.1807** | **5.263** |
| 2 | 3 | 1 | 0.006411 | 0.1135 | 5.652 |
| 3 | 5 | 56 | 0.002229 | 0.1287 | 5.748 |

H

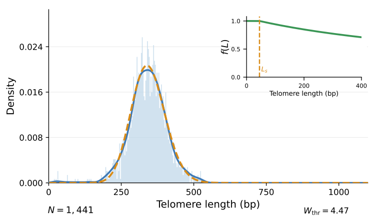


| rank | $a$ | $L_s$ | $\beta$ | $p$ | $W_1$ |
|---|---|---|---|---|---|
| **1** | **4** | **45** | **0.001147** | **0.1867** | **3.224** |
| 2 | 5 | 163 | 0.001049 | 0.1686 | 3.295 |
| 3 | 6 | 183 | 0.001424 | 0.1362 | 3.447 |

I

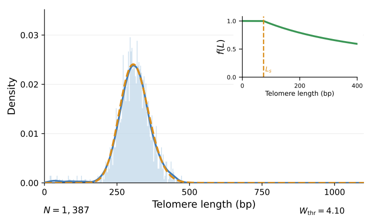


| rank | $a$ | $L_s$ | $\beta$ | $p$ | $W_1$ |
|---|---|---|---|---|---|
| **1** | **4** | **74** | **0.002108** | **0.1674** | **3.289** |
| 2 | 3 | 21 | 0.004342 | 0.1488 | 3.342 |
| 3 | 2 | 2 | 0.008691 | 0.1374 | 3.398 |

**Supplementary Figure S14. Wild-type W303 chromosome-end-specific telomere length distributions fitted by our model (Part 2). (A–I)** Same as Supplementary Fig. S13, for the indicated chromosome ends.

**A**

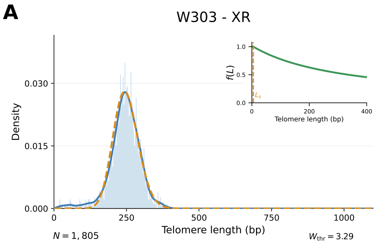


| rank | $a$ | $L_s$ | $\beta$ | $p$ | $W_1$ |
|---|---|---|---|---|---|
| **1** | **3** | **5** | **0.003018** | **0.1934** | **5.015** |
| 2 | 2 | 0 | 0.009971 | 0.1461 | 5.308 |
| 3 | 4 | 54 | 0.002734 | 0.1645 | 5.447 |

**B**

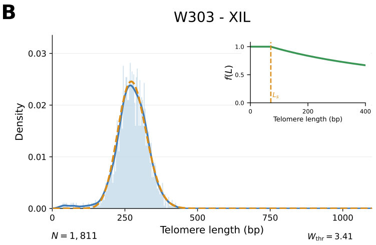


| rank | $a$ | $L_s$ | $\beta$ | $p$ | $W_1$ |
|---|---|---|---|---|---|
| **1** | **4** | **71** | **0.001506** | **0.1915** | **3.619** |
| 2 | 3 | 2 | 0.004397 | 0.1524 | 3.891 |
| 3 | 2 | 0 | 0.010576 | 0.1294 | 4.152 |

**C**

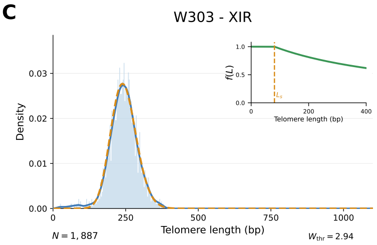


| rank | $a$ | $L_s$ | $\beta$ | $p$ | $W_1$ |
|---|---|---|---|---|---|
| **1** | **4** | **81** | **0.001932** | **0.1910** | **2.699** |
| 2 | 3 | 45 | 0.007160 | 0.1394 | 3.253 |
| 3 | 2 | 62 | 0.020848 | 0.1077 | 3.485 |

**D**

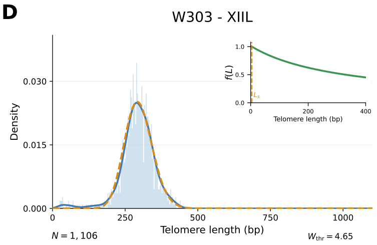


| rank | $a$ | $L_s$ | $\beta$ | $p$ | $W_1$ |
|---|---|---|---|---|---|
| **1** | **3** | **4** | **0.003072** | **0.1766** | **5.315** |
| 2 | 2 | 22 | 0.009787 | 0.1377 | 5.515 |
| 3 | 4 | 54 | 0.002412 | 0.1588 | 6.264 |

**E**

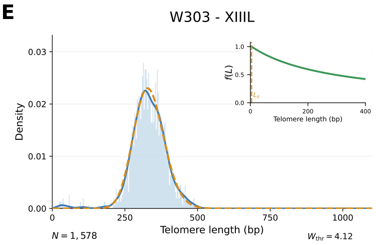


| rank | $a$ | $L_s$ | $\beta$ | $p$ | $W_1$ |
|---|---|---|---|---|---|
| **1** | **3** | **4** | **0.003464** | **0.1567** | **3.669** |
| 2 | 4 | 72 | 0.002542 | 0.1511 | 3.674 |
| 3 | 2 | 17 | 0.009522 | 0.1264 | 3.688 |

**F**

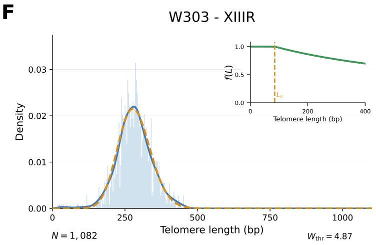


| rank | $a$ | $L_s$ | $\beta$ | $p$ | $W_1$ |
|---|---|---|---|---|---|
| **1** | **5** | **85** | **0.001372** | **0.1578** | **2.881** |
| 2 | 4 | 30 | 0.000998 | 0.2000 | 2.896 |
| 3 | 6 | 118 | 0.001792 | 0.1293 | 2.916 |

**G**

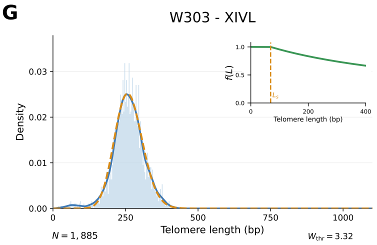


| rank | $a$ | $L_s$ | $\beta$ | $p$ | $W_1$ |
|---|---|---|---|---|---|
| **1** | **4** | **69** | **0.001521** | **0.1939** | **3.151** |
| 2 | 3 | 11 | 0.005905 | 0.1360 | 3.222 |
| 3 | 2 | 0 | 0.011702 | 0.1252 | 3.272 |

**H**

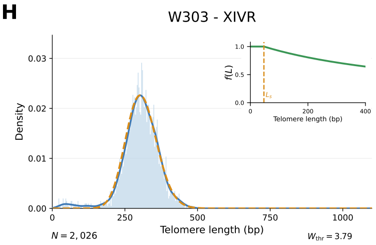


| rank | $a$ | $L_s$ | $\beta$ | $p$ | $W_1$ |
|---|---|---|---|---|---|
| **1** | **4** | **47** | **0.001570** | **0.1780** | **5.243** |
| 2 | 3 | 0 | 0.004617 | 0.1390 | 5.447 |
| 3 | 2 | 0 | 0.010196 | 0.1227 | 5.631 |

**I**

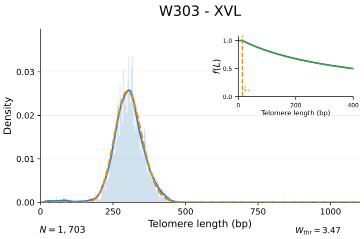


| rank | $a$ | $L_s$ | $\beta$ | $p$ | $W_1$ |
|---|---|---|---|---|---|
| **1** | **3** | **14** | **0.002582** | **0.1913** | **3.275** |
| 2 | 4 | 89 | 0.002511 | 0.1630 | 3.298 |
| 3 | 2 | 1 | 0.007747 | 0.1507 | 3.298 |

**Supplementary Figure S15. Wild-type W303 chromosome-end-specific telomere length distributions fitted by our model (Part 3). (A–I)** Same as Supplementary Fig. S13, for the indicated chromosome ends.

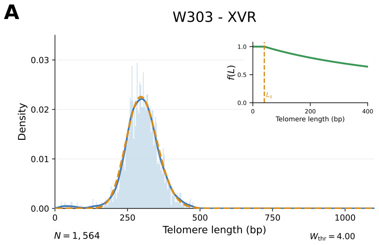


| rank | $a$ | $L_s$ | $\beta$ | $p$ | $W_1$ |
|---|---|---|---|---|---|
| **1** | **4** | **40** | **0.001545** | **0.1787** | **3.128** |
| 2 | 3 | 0 | 0.005259 | 0.1304 | 3.190 |
| 3 | 2 | 8 | 0.011865 | 0.1136 | 3.275 |

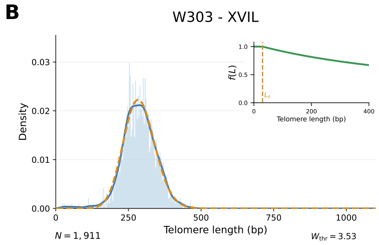


| rank | $a$ | $L_s$ | $\beta$ | $p$ | $W_1$ |
|---|---|---|---|---|---|
| **1** | **4** | **30** | **0.001331** | **0.1866** | **2.591** |
| 2 | 5 | 109 | 0.001423 | 0.1599 | 2.690 |
| 3 | 3 | 7 | 0.006802 | 0.1160 | 2.937 |

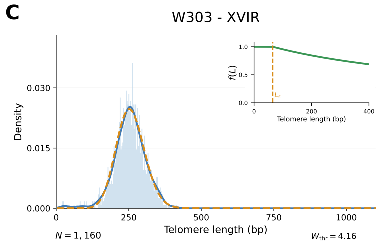


| rank | $a$ | $L_s$ | $\beta$ | $p$ | $W_1$ |
|---|---|---|---|---|---|
| **1** | **4** | **65** | **0.001358** | **0.1990** | **3.170** |
| 2 | 3 | 2 | 0.005962 | 0.1341 | 3.435 |
| 3 | 5 | 78 | 0.002899 | 0.1330 | 3.530 |

**Supplementary Figure S16. Wild-type W303 chromosome-end-specific telomere length distributions fitted by our model (Part 4). (A–I)** Same as Supplementary Fig. S13, for the indicated chromosome ends.

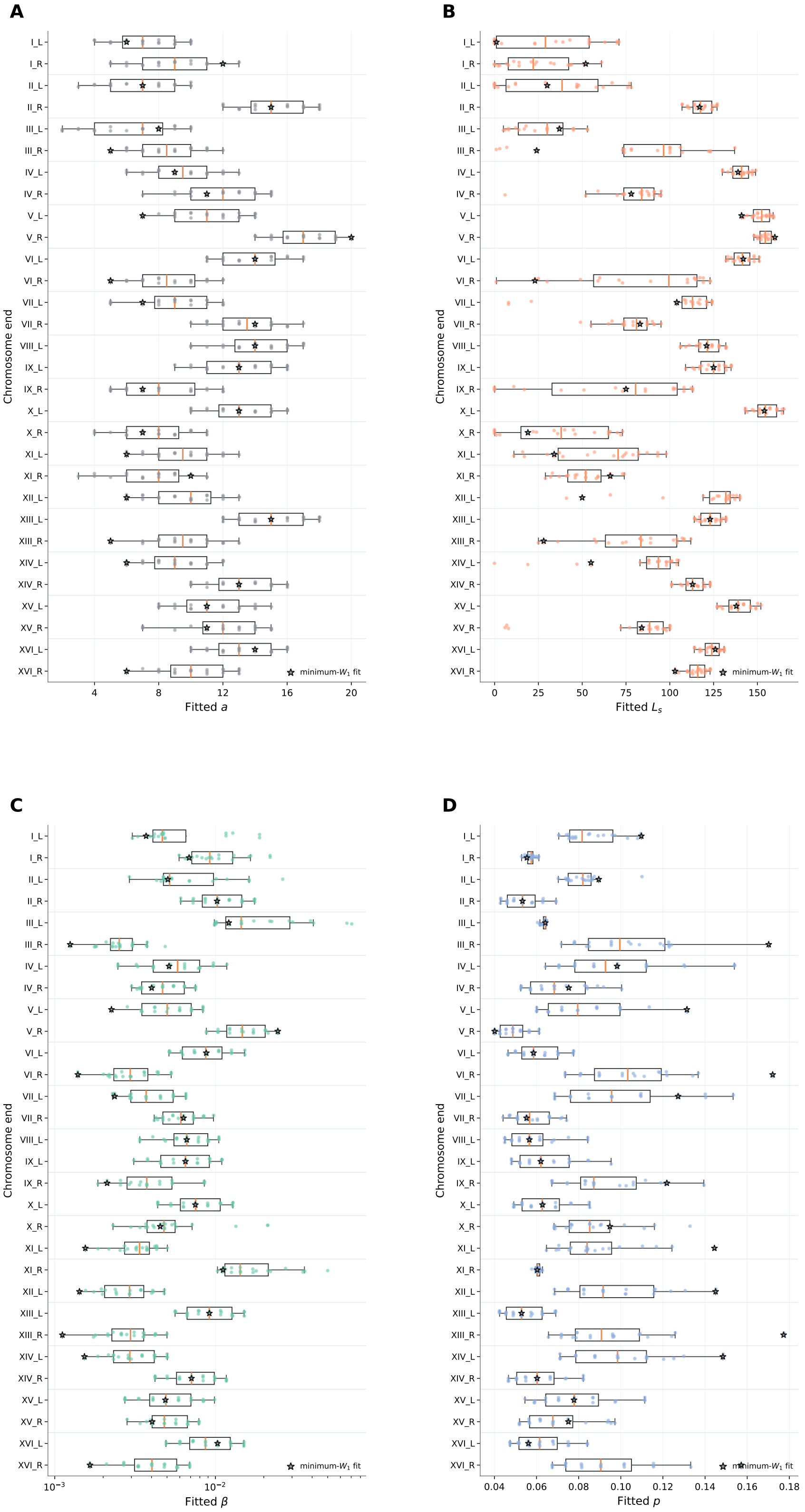


**Supplementary Figure S17. Chromosome-end-specific variability in fitted parameters for pooled data.** Measurements from three *yku70Δ* replicates were pooled and analyzed separately for each of the 32 chromosome ends. CMA-ES optimization yielded acceptable fits for 30 chromosome ends. Boxplots show the distributions of fitted parameters across optimization runs: **(A)** $a$, **(B)** $L_s$, **(C)** $\beta$, and **(D)** $p$. Boxes indicate the interquartile range, central lines the median, and whiskers the spread of fitted values. The best-fit parameter is indicated by a star. The parameter $\beta$ is shown on a logarithmic scale.
